# A multimodal, correlative magnetic tweezers–TIRF platform for high-throughput single-molecule interrogations

**DOI:** 10.64898/2026.08.11.744120

**Authors:** M. Sadegh Feiz, Jelmer Cnossen, Yibulayin Wubulikasimu, Salina Quack, Thomas Bugea, Andrei Zupnik, Ranjit K. Prajapati, Asif Rakib, Flavia S. Papini, Quinte Smitskamp, Anssi M. Malinen, David Dulin

## Abstract

Single-molecule techniques can resolve biological reactions at unmatched detail, but their low throughput and single-modality readouts have kept them out of data-intensive pipelines such as omics and drug discovery, and beyond reach of low-yield biological systems. Here we introduce a multimodal platform integrating high-throughput magnetic tweezers with ultra-wide-and flat-field objective-based total internal reflection fluorescence, enabling simultaneous force, torque, multicolor fluorescence, and temperature-dependent measurements on up to thousands of individual molecules in parallel and in real time. We demonstrate accurate single-molecule Förster resonance energy transfer (smFRET) for prism-based spectral imaging, capture temperature-dependent hairpin folding dynamics at high temporal resolution with smFRET and use correlative torque–fluorescence measurements to unravel the open-complex formation dynamics during bacterial transcription initiation. By unifying high resolution, throughput, and multimodal readout, this platform enables multidimensional dissection of complex biomolecular reactions with high statistical confidence, unlocking single-molecule biophysics for integration with drug discovery, omics, and cryo-EM workflows.

## Introduction

Biological reactions at the molecular scale proceed through multiple, interconnected pathways composed of consecutive, often reversible, stochastic steps, giving rise to highly heterogeneous dynamics (*1*). Single-molecule biophysics techniques are uniquely suited to disentangle this heterogeneity by tracking individual molecules throughout the reaction (*2, 3*). Yet capturing these reactions in their full complexity — including the rarest events — requires extensive statistics across many experimental conditions and modalities, together with the millisecond temporal and nanometer spatial resolution needed to follow them (*4–6*). Meeting these requirements is central to integrating single-molecule approaches into structural biology (*7, 8*) and omics (*6*) workflows, where they can provide a dynamic link between structure, sequence, and function. To date, however, multimodal single-molecule assays deliver either high spatiotemporal resolution at low throughput — as with optical tweezers combined with fluorescence microscopy (*9, 10*) — or the converse, as with flow stretching (*11*), leaving a clear technological gap.

Magnetic tweezers (MT) and total internal reflection fluorescence (TIRF) microscopy are particularly well suited to bridge this gap. MT applies force or torque to a biomolecule tethered to the surface of a flow chamber via a magnetic bead, enabling parallel interrogation of hundreds of biomolecular reactions at nanometer and millisecond resolution over a force range spanning femtonewtons to nanonewtons (*12–14*). This versatility has driven major advances in our understanding of the mechanical properties of nucleic acids and proteins (*15, 16*), the dynamics of nucleic-acid-processing enzymes (*17–21*), and the mechanism of action of small-molecule drugs targeting them (*22–24*). TIRF microscopy, in turn, is a surface-based wide-field fluorescence technique capable of imaging hundreds to thousands of fluorescently labeled biomolecules across multiple spectral channels (*25–28*), and has been used extensively to track biomolecular complex formation (*29*) and conformational dynamics through smFRET (*30, 31*). Both MT and TIRF are surface-based, making them natural candidates for correlative single-molecule measurements; however, only low-throughput implementations have been reported to date (*32–40*). While wide-field multispectral single-molecule TIRF with homogeneous illumination has been demonstrated using prism-based (*27, 29*) and waveguide-based (*41, 42*) geometries, achieving comparable performance in objective-based TIRF has remained particularly challenging. This bottleneck has constrained integration with other high-throughput technologies — including magnetic tweezers — and limits the possible throughput when single-molecule TIRF is combined with other modalities, such as next generation sequencing (*6*).

Here, we introduce a high-throughput, correlative MT–TIRF platform that performs simultaneous force, torque, multicolor fluorescence, and temperature-dependent measurements on up to thousands of single molecules in parallel. To maximize throughput while minimizing the number of scientific CMOS (sCMOS) cameras required for fluorescence imaging, we developed a super wide field homogenous illumination scheme for objective-based TIRF that covers the full sensor area of a single sCMOS camera and a prism-based two-color spectral imaging system. Together, they enable simultaneous monitoring of thousands of fluorescently labeled molecules on a single detector. We first benchmarked the TIRF modality for accurate smFRET (*43*) and applied it to resolve the temperature dependence of hairpin folding-unfolding dynamics at high temporal resolution. In correlative mode, the instrument simultaneously interrogates up to 400 tethers in real time with combined force, torque, and fluorescence readout. To illustrate this capability, we examined open-complex formation during bacterial transcription initiation, using the TIRF channel to monitor binding of the initiation complex and MT torque spectroscopy to correlate binding with promoter opening to show that open-complex formation is rate-limited by a single kinetic transition.

By unifying high resolution, high throughput, and multimodal readout in a single instrument, this platform opens the way to high-precision, data-driven dissection of complex stochastic processes and their integration with structural and omics datasets.

## Results

### A microscope design for high-throughput MT and TIRF measurements

High-throughput MT–TIRF demands a large field of view in both modalities. We extended our previous high-throughput MT design (*14, 44, 45*) into a bespoke inverted microscope combining MT imaging, TIRF laser excitation, and fluorescence detection over a shared large field of view (**Fig. 1**, **Fig. S1a**). A ∼1 nm-resolution autofocus referenced to a surface-immobilized bead maintained long-term focal stability, matching the performance of our previous design (*46*) (<0.02 nm at 10 s integration; **Fig. S1c**, **Supplementary Information**). Magnet positioning and high-power LED illumination followed previously described designs (*14, 45*). Temperature was regulated to 0.1 °C with a resistive foil heater on the objective (*47*).

**Figure 1:**
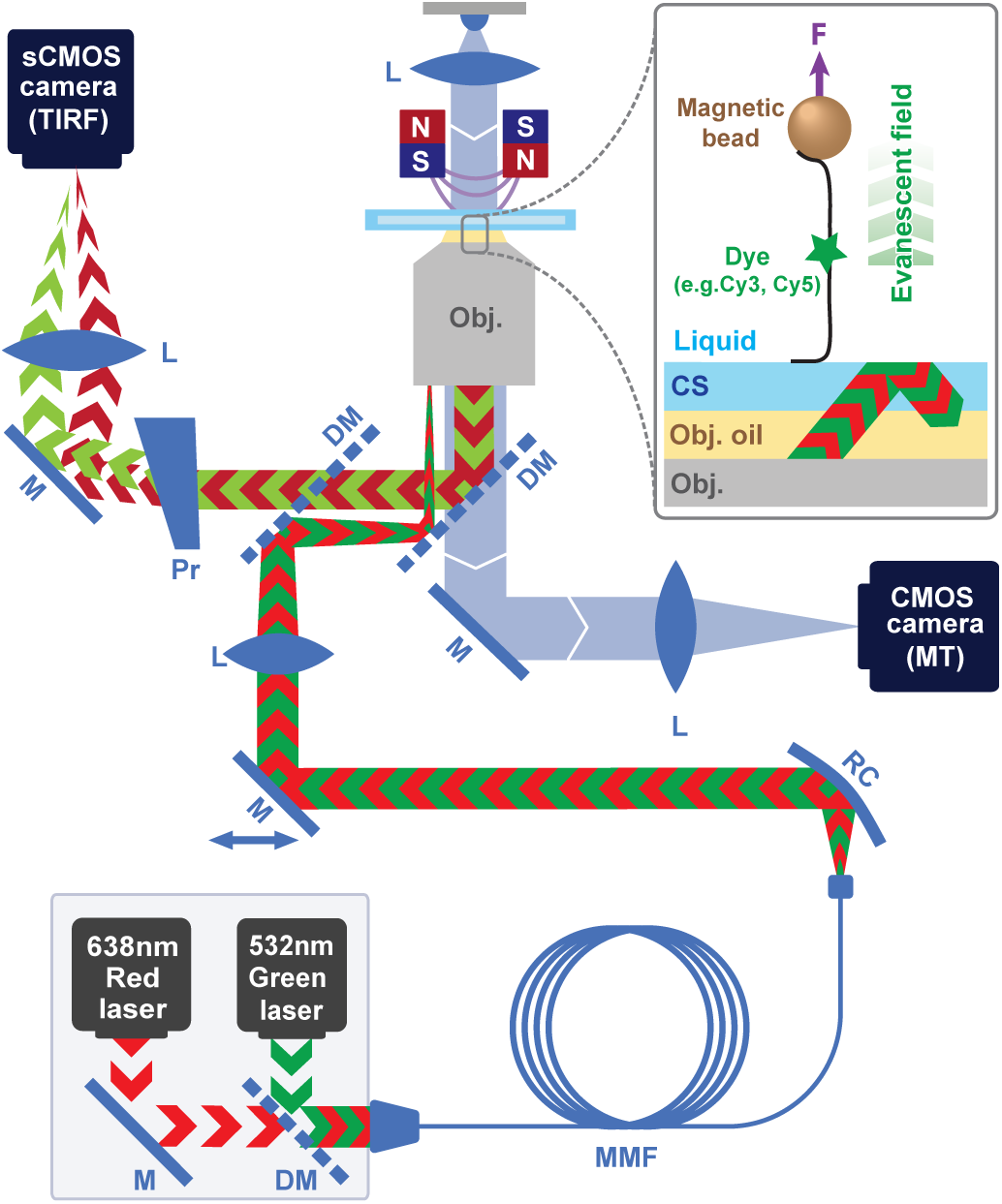
Schematic description of the high-throughput multimodal MT-TIRF assay. The schematic represents the design for objective-based TIRF for large field of view illumination, wedge prism-based multispectral imaging, and MT illumination and imaging. L, M, DM, Pr, RC, MMF and Obj stand for lens, mirror, dichroic mirror, prism, reflective collimator, multi-mode fiber, and objective, respectively. Inset: nucleic acid tethered magnetic bead.

We next homogenized the illumination across the sCMOS sensor. The excitation laser was coupled into a multimode fiber (MMF) coiled and vibrated by a piezoelectric disc and a mechanical vibrator, scrambling the propagating modes and lowering speckle contrast (*48*) (**Fig. S2ab**, **Supplementary Information**). At 532 nm, the contrast minimized at the ∼1.3 kHz piezo resonance, while at 638 nm excitation, which showed intrinsically low contrast, the contrast was largely unaffected by the speckle-reduction unit (**Fig. S2cde**). The resulting illumination was uniform across a (210 x 210) µm^2^ field of view spanning the full sCMOS sensor, yielding homogeneous emission from both fluorescent beads (**Fig. S2fgij**, **Methods**) and single fluorophores (**Fig. S3a–d**).

We calibrated the evanescent field depth from the autofluorescence of MyOne magnetic beads tethered by a torsionally constrained double-stranded RNA (dsRNA) (**Methods, Supplementary Information**). Rotating the magnets progressively supercoiled the dsRNA, drawing the bead toward the flow-chamber surface (**Fig. S3e**). The penetration depth was ∼90 nm (**Fig. S3f**), consistent with previous reports (*49, 50*). Autofluorescence vanished beyond 500 nm from the surface, confirming effective total internal reflection.

### Establishing prism-based multispectral imaging

Our integrated platform enables force, torque, fluorescence, and correlative measurements, and high-throughput capability is empowered by a large chip area CMOS camera. In MT and TIRF experiments, raw images are segmented into regions of interest (ROIs) around a magnetic bead or a fluorescent emitter, respectively. MT ROIs are ∼10 µm wide (*12, 14*), whereas fluorescence ROIs are <1 µm for diffraction-limited emitters (*51*). Maximizing correlative throughput therefore requires packing as many MT ROIs as possible into a single field of view. To keep the optics simple — two cameras (MT and TIRF) with minimal imaging-path components — we used prism-based spectral separation (*52–56*) to spatially offset the point spread functions of multiple fluorophores (here Cy3B and ATTO647N) onto a single sCMOS camera (**Fig. 1**, **Fig. S1a**, **Fig. S4ab**). Fluorescent bead imaging yielded a 9.6 ± 0.6 pixel separation between the 532 nm and 638 nm channels, in agreement with our calculations (**Fig. S4cd**, **Table S1**, **Supplementary Information**), with ∼1 pixel dispersion across the field of view (**Fig. S4de**). The strategy is easily extendable to a broad range of emission wavelengths, i.e. from 500 to 800 nm (**Fig. S4b**), with negligible impact on throughput, enabling high-throughput objective-based TIRF over a broad spectral range.

### High-throughput single-molecule FRET

Single-molecule FRET (smFRET) measures conformational dynamics in biomolecules with near-structural resolution (*43, 57, 58*), greatly complementing static structural methods, such as cryo-EM. To benchmark the assay, we designed two double-stranded DNA FRET standards labeled with Cy3B (donor) and ATTO647N (acceptor), with the fluorophores separated by 11 or 27 base pairs and predicted FRET efficiencies of *E*_expected_ ≈ 0.77 (high-FRET) and *E*_expected_ ≈ 0.10 (low-FRET) (*59*). We used alternating laser excitation (ALEX) with stoichiometry–efficiency (S–E) analysis to correct the FRET signals (*60*) (**Fig. 2ab**, **Fig. S5abc**, **Methods**), recording donor-only, acceptor-only, and dual-labeled constructs in ALEX mode to extract the correction parameters (**Fig. 2a**; **Supplementary Information**). The high-throughput TIRF configuration simultaneously detected nearly 4,000 single DNA molecules in a single field of view (**Fig. 2a**). After correcting the raw S–E distributions (**Methods**, **Supplementary Information**, **Fig. S5ab**) (*43*), the low-and high-FRET constructs gave corrected efficiencies of *E*_corr_ = 0.11 ± 0.07 and *E*_corr_ = 0.74 ± 0.06 (mean ± s.d.), respectively, in close agreement with predictions.

**Figure 2:**
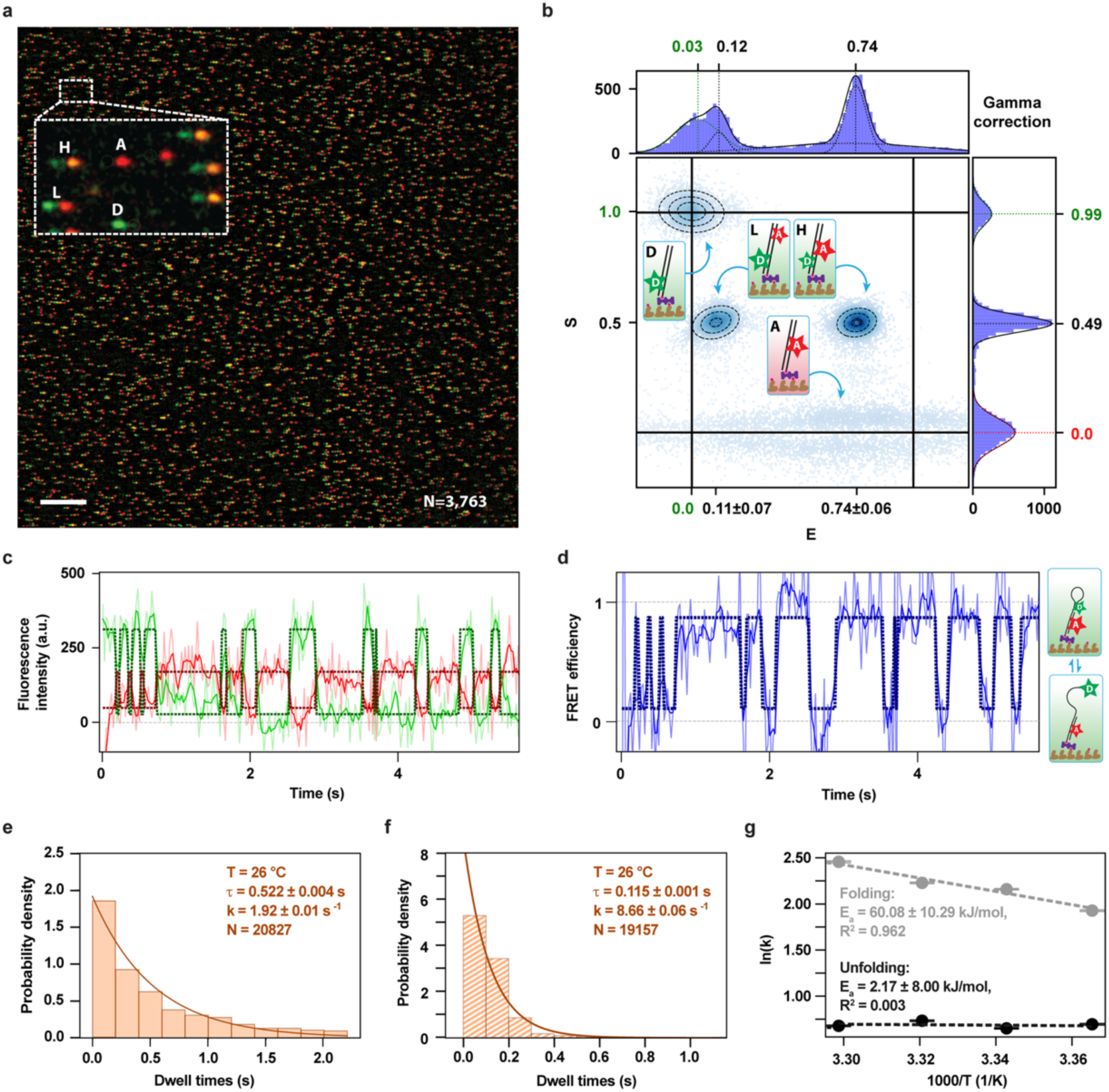
Wide-and flat-field TIRF for high-throughput accurate and dynamic smFRET. **(a)** TIRF camera Field of view The Cy3B and ATTO647N labeled oligos. Inset: zoom-in of the field of view showing low-FRET (L), high-FRET (H), donor-only (D), and acceptor-only (A) populations. 532 nm and 638 nm laser excitations are represented in green and red, respectively, and FRET in yellow. The scale bar indicates 20 µm and the total number of detected fluorescent emitters is N=3,763. **(b)** Corrected S-E plot for the data acquired as described in (a). The dashed oval lines represent 2D Gaussian fits. E and S histograms and their respective 1D-Gaussian fit (solid lines) are represented above and on the right of the S-E plot. The values indicated in green and red correspond to the D-only and A-only populations, respectively. The schematics represent the different duplex designs with the acceptor (A) and donor dye (D), while the black characters report on the populations described in (a). The errors for E values are calculated from the 2D Gaussian fit, showing the standard deviation alongside the E axis. **(c)** Acceptor and donor intensity and **(d)** FRET efficiency as a function of time for a DNA hairpin with a 6 bp stem and 30 nt loop alternating between a folded (high FRET) and unfolded (low FRET) state at 26°C (**Methods**). The full trace is shown in **Figure S6c**. The light and dark color are the raw and 4-frame averaged traces. The dashed line is the HMM fit to the raw data. **(e, f)** Dwell time histograms of the unfolding (e) and folding (f) dynamics at 26°C. A mono-exponential fit is shown as a solid line and the mean dwell time ± standard error of the mean, the reaction rate k and the number of state detection n are given in the plot. **(g)** Arrhenius plot of the of unfolding (black, kunfolding) and folding (grey, kfolding) rates as a function of temperature. The Arrhenius fit is shown as dashed line (**Eq. 45**). The activation energy Ea and the coefficient of determination R² are indicated in the plot. The error bar was estimated through linear error propagation.

We next used this smFRET assay to monitor folding–unfolding dynamics of a DNA hairpin as a function of temperature (*47*). We assembled a hairpin with a short stem (6 bp) and long polyadenosine loop (32 nt) that can be in either a folded or unfolded state (**Methods**), with Cy3B (donor) and ATTO647N (acceptor) labeling on opposite strands (**Fig. 2c-g**, **Fig. S6**, **Supplementary Information**) (*47, 61, 62*). We extracted the unfolded–folded dynamics with hidden Markov modeling (HMM) (**Fig. 2cd**, **Fig. S6a**), recovering temperature-independent corrected FRET efficiencies of ∼0.09 and ∼0.85 for the unfolded and folded state, respectively (**Fig. S6b**) (*61*). Folding and unfolding kinetics were well described by single exponentials (**Fig. 2ef**, **Fig. S6c**). The folding rate increased from 6.9 ± 0.1 s⁻¹ at 24°C to 11.7 ± 0.1 s⁻¹ at 30°C (**Fig. S6c**), yielding an apparent activation energy *E_A_* = 60 ± 10 kJ/mol (*63, 64*). The unfolding rate was temperature-independent (2.00 ± 0.02 s⁻¹ and 1.97 ± 0.02 s⁻¹ at 24°C and 30°C; (*E_A_* = 2.17 ± 8.00 kJ/mol)), consistent with the high GC content (83%) of the duplex (*65, 66*). Together, these experiments establish our TIRF assay as a platform for accurate smFRET measurements of biomolecular dynamics at 40 Hz image acquisition and across temperature, on thousands of molecules in parallel.

### Mapping biomolecules positions on both MT and TIRF cameras

Correlative measurements require coordinate mapping between the MT and TIRF cameras so that the measured tether attachment position predicts the fluorescence signal position on the TIRF sCMOS. We performed this mapping in two steps (**Methods**). First, we imaged and tracked 500 nm fluorescent beads on both cameras to compute a transfer matrix correlating ROIs across the two fields of view (**Methods**). Second, we corrected for off-axis attachment of the biomolecule to the magnetic bead: because the MT tracker locates the bead center, the apparent tether position can deviate from the true attachment site by up to the bead radius, on the order of microns (*67–69*) (**Methods**). We recovered the true attachment position by rotating the magnets and centering the resulting circular bead trajectory (**Fig. S7a**, **Fig. 3a**) (*69*). Deviations from parallel field geometry produce more complex trajectories, described elsewhere (*69*) (**Fig. S7b**); these were rare in our configuration and were discarded from subsequent analysis.

**Figure 3:**
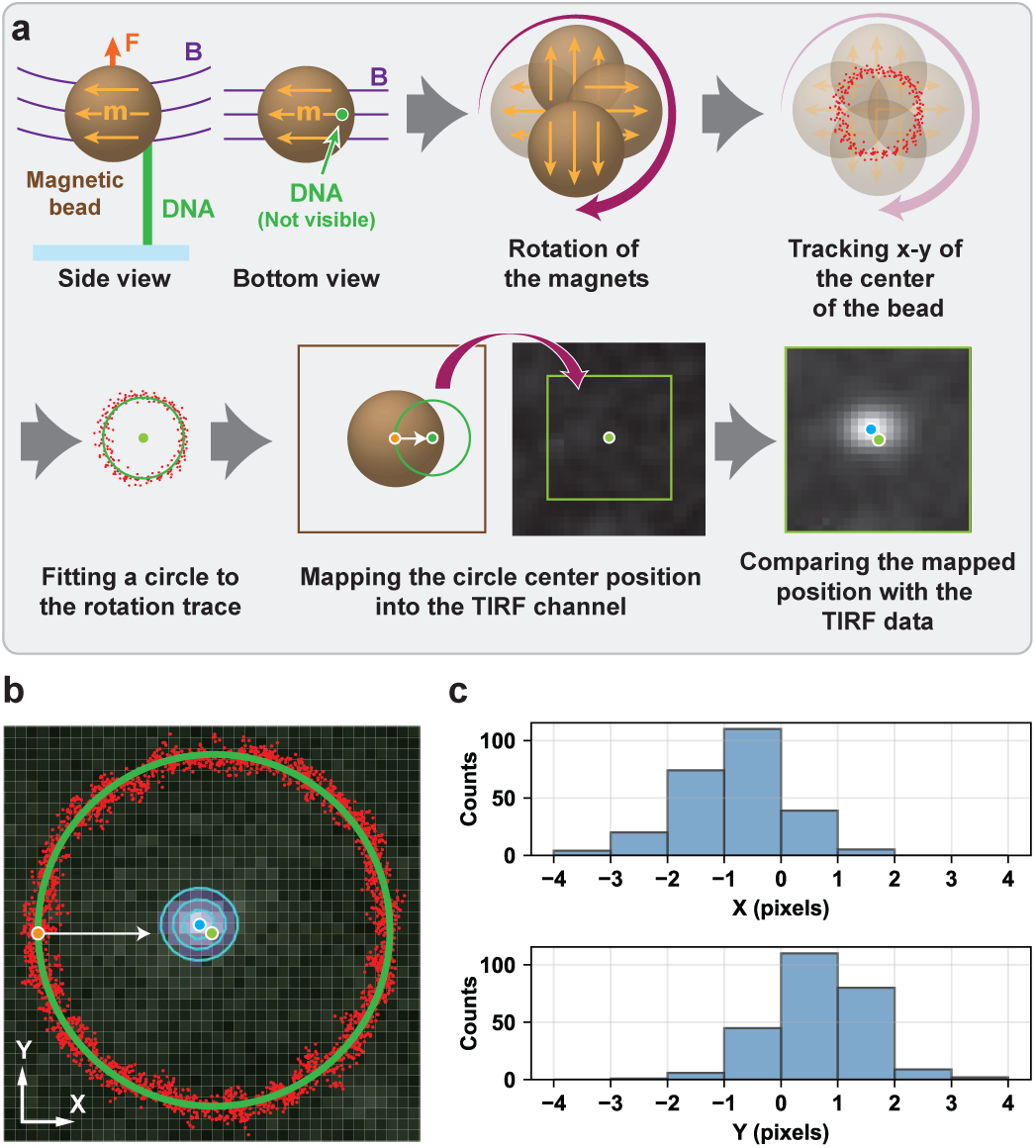
Absolute tether positioning on both the MT and TIRF cameras. **(a)** Schematic of the procedure to define the attachment position of a tether on the flow cell surface for both the MT and TIRF camera. ***m*** is the magnetic bead magnetization, ***B*** is the magnetic field and ***F*** is the force. **(b)** Red dots: y vs. x position of the magnetic bead when rotating the magnets ±1 turns at 0.02 turns per second and with *F* = 5 *pN*. The solid green circle is a *r* = +*x*2 + *y*2 fit to the data. The horizontal and vertical grey grid is the MT camera pixel mapping on the sample plane, with the separation of ∼0.106 µm. The green dot indicates the center of the fitted circle, which is supposed to be the attachment position of the tether. The MT trace is overlapped on the TIRF camera image. The pixels of the TIRF camera are separated by ∼0.107 µm in the sample plane. The blue contours represent a 2D Gaussian fitting on the recorded TIRF image and hence the blue dot shows the tether to surface attachment position. The orange dot shows the initial bead center position. **(c)** Histogram of the x-and y-axis offset calibration for measurement performed as described in (a and b) for N=252 tethers.

We validated the mapping by directly visualizing the nucleic acid tether with the fluorescent intercalator SYTOX Orange (**Methods**, **Fig. 3b**). The imaged attachment position deviated from the rotation-derived estimate by a small systematic offset of (−0.86 ± 0.06) *pixels* and (0.68 ± 0.05) *pixels* (mean ± s.e.m.) along x and y, respectively (**Fig. 3c**), which we incorporated into subsequent tether localization. This label-free method, implemented within a magnetic tweezers instrument, determines tether attachment positions across multiple detectors with sub-pixel precision (patent pending, EP25186932).

### High-throughput correlative MT–TIRF

Next, we integrated high-throughput MT with TIRF microscopy for parallel correlative single-molecule measurements. Common magnetic beads (e.g., MyOne) are strongly autofluorescent (**Fig. S3ef**), precluding single-dye detection at short tethers or low forces. We identified two brands with minimal autofluorescence (**Fig. S8a**) and acceptable force dispersion (**Fig. S8bc**): 1 µm NEB beads for low-force and 3.3 µm Compel beads for high-force measurements (**Supplementary Information**).

To benchmark the throughput capabilities of the correlative MT–TIRF platform, we monitored extension of coilable ∼3.2 kbp dsRNA tethers while injecting the fluorescent intercalator SYBR™ Gold (**Methods**). Because SYBR™ Gold unwinds duplex nucleic acids upon intercalation (*70*) (**Fig. 4a**), we expected plectoneme formation at low force. Tracking ∼400 tethers simultaneously in both channels at 0.3 pN (**Fig. 4bc**), we found that SYBR™ Gold rapidly induced plectonemes in the coilable subpopulation (one-third of tethers (*71*)) — a drop in tether extension with a concurrent TIRF fluorescence rise as plectonemes entered the evanescent field — while non-coilable molecules retained stable extension and constant fluorescence. Prolonged excitation then caused SYBR™ Gold-mediated dsRNA photodamage (*72*), nicking the backbone and abolishing coilability; the resulting supercoil relaxation produced extension recovery (**Fig. 4de**). These results establish parallel, high-throughput force, torque, fluorescence, and correlative interrogation across hundreds of single nucleic acid tethers.

**Figure 4.**
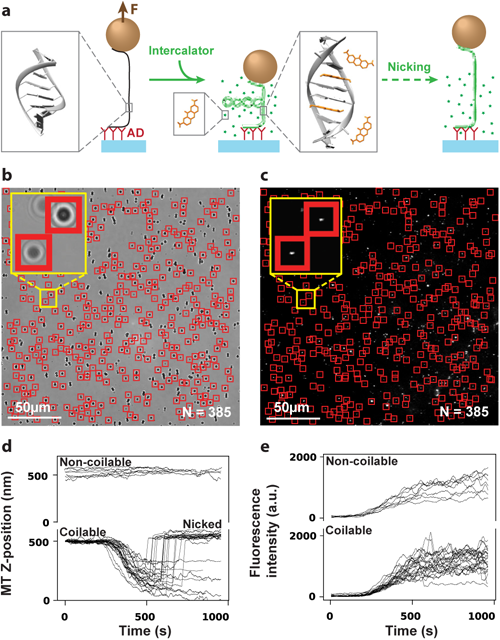
Correlative assessment of the MT and TIRF channels. **(a)** The schematic describing Binding fluorescent intercalators (SYBR™ Gold) on dsRNA causing twist and potentially inducing nicks. The structures are adapted from PDB 2KD4. **(b, c)** Images of a same field of view from the camera of the MT and TIRF channels, respectively. Nucleic acid tether is fluorescently labelled with SYBR™ Gold intercalator. **(d)** Time traces of the magnetic bead z-position for coilable and non-coilable tethers upon addition of SYBR™ Gold. **(e)** Correlated fluorescence intensity for the same tethers as in (d).

### Capturing the bacterial RNA polymerase open complex formation in real-time

We next applied the platform to interrogate a key regulatory step in bacterial transcription: conversion of the RNA polymerase (RNAP)–promoter closed complex (RPC) into the open complex (RPO). In *Escherichia coli*, the RNAP core enzyme associates with the housekeeping initiation factor σ^70^ to form the holoenzyme, which binds promoter DNA to form the RPC and then melts the promoter to form the RPO, opening the transcription bubble for RNA synthesis (*73, 74*). Structural intermediates along this pathway have been resolved by cryo-EM (*75, 76*) and kinetic schemes refined by biochemical analyses (*74, 77*), yet single-molecule readouts remain partial: MT alone resolve bubble dynamics but cannot monitor holoenzyme binding (*78, 79*); single-molecule fluorescence microscopy visualizes binding without reporting on bubble formation (*80, 81*); and a recent fluorescence enhancement assay captures holoenzyme association and initial promoter unwinding simultaneously but without a direct readout of bubble size (*82*). A correlative assay combining these orthogonal modalities would therefore complement existing approaches and enable investigation of transcription initiation downstream of bubble formation.

We designed a coilable DNA construct encoding the consensus lac promoter lacCONS+2 (**Methods**) (*79, 83*). The promoter was positioned adjacent to the digoxigenin-labeled handle anchoring the construct to the flow-chamber surface (**Fig. 5a**), maximizing the TIRF fluorescence intensity of the ATTO550-labeled holoenzyme upon promoter binding (**Methods**). Magnetic tweezers, owing to their high sensitivity to DNA twist change under supercoiling (*12*), precisely detect transcription bubble formation and dynamics (*78, 79, 84*). Each tether was held at ∼0.3 pN and negatively supercoiled by −4 turns to generate plectonemes (*15*) (**Fig. 5a**). Injection of labeled holoenzyme produced discrete TIRF fluorescence signals at the promoter region, indicating RPC formation (**Fig. 5bc**). The magnetic bead then moved upward by 40 ± 16 nm (mean ± s.d.; **Fig. S9ab**), consistent with reduced writhe upon promoter melting and RPO formation (**Fig. 5ac**) (*78, 79*). The RPO lifetime (hundreds of seconds (*79*)) substantially exceeded the fluorophore photobleaching lifetime (tens of seconds), so each molecule contributed a single opening event. The MT–TIRF assay enables the collection of a large dataset of RPC-to-RPO transition time (N = 120) (**Fig. S9a**, **Fig. 5d**). The resulting distribution was well described by a single exponential with exit rate *k_RP_*C = (0.069 ± 0.006) *s*^−1^ (mean ± s.d.), indicating that a single long-lived state dominates the RPC-to-RPO transition.

**Figure 5:**
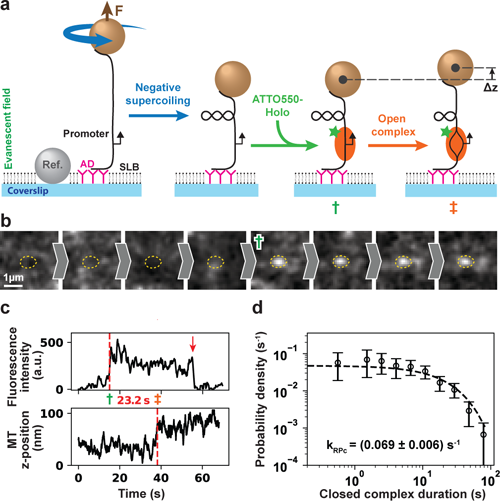
Monitoring bacterial RNA polymerase open-complex formation using high-throughput correlative MT-TIRF. **(a)** Schematic of the MT-TIRF experiment. The binding of the ATTO550-labeled holoenzyme (marked by green †) is detected using TIRF, while the open complex formation (marked by orange ‡) is monitored using negatively supercoiled DNA with MT. **(b)** TIRF region of interest reporting holoenzyme binding (green †) at the predicted promoter position (yellow oval). **(c)** Correlative-time traces of fluorescent intensity and magnetic bead z-position vs. time for the experiment described in (a) and the region of interest represented in (b). The red dashed lines highlight the fluorescence intensity increase time from the holoenzyme binding (green †) and the magnetic bead z-position change from the promoter opening (orange ‡). The time delay is indicated between the two red dashed lines. **(d)** The probability density distribution of the time delay between holoenzyme binding and promoter opening (N=120). The error bars represent 95% confidence intervals from 1000 bootstrap samples. The error estimate for the rate is the standard deviation obtained through a separate set of 100 bootstrap samples.

Our measurements are broadly consistent with recent single-molecule fluorescence studies (*82, 85*), though with subtle differences. Mazumder and colleagues reported a peaked dwell-time distribution for initial promoter unwinding, suggesting two sequential steps of comparable lifetime (*82*), whereas our data follow single-exponential kinetics (**Fig. 5d**) (similarly to Malinen and colleagues, though they reported faster kinetics (*85*)). Whether the MT signal here and the fluorescence enhancement of ref. (*82*) report on the same rate-limiting steps will require further investigation. Small differences in the experiments design can explain these discrepancies, such as the construct length upstream the-35 region (which interacts with RNAP α-CTD and impacts transcription initiation (*85*)), temperature at which experiments were performed (which affects RPO dynamics (*79*)), negatively supercoiled promoter or buffer composition (also affecting RPO dynamics (*78, 79*)). Last, site-specific ATTO550 labeling of σ^70^ by mutating two of the three cysteine residues could have also affected the kinetics (**Methods**). However, the exit rate measured here agrees with our previous MT-based estimate using wild-type σ^70^ (*79*), supporting the present measurement (**Fig. 5d**). Future implementations will combine orthogonal distance readouts, such as smFRET (**Fig. 2**), with enhanced MT temporal resolution to resolve short-lived intermediates in transcription bubble formation, as predicted by cryo-EM (*75, 86*).

## Discussion

Single-molecule biophysics has transformed our understanding of biological systems through direct manipulation and visualization of individual biomolecules while performing their activity (*3*). Throughput has long been the central constraint, restricting access to intricate reaction pathways and to complex, low-yield reactions such as the assembly of multimeric protein complexes with diverse cofactors (*4, 5*). The limitation is particularly acute for correlative assays combining force, torque, and fluorescence spectroscopy (*12, 87*). As technologies that interrogate heterogeneity at unprecedented scale reach maturity — cryo-EM for biomolecular complex organization and composition, and next-generation sequencing for nucleic acid sequence — single-molecule methods must match this throughput to map structure–function and sequence–function relationships.

Large-area, high-speed CMOS and sCMOS cameras, together with massively parallel computation on graphics processing units (GPUs), have made high-throughput single-molecule acquisition and real-time analysis accessible. High-throughput implementations have accordingly emerged for individual modalities, including TIRF microscopy (*27, 88*), MT (*46, 89, 90*), tethered-particle motion (*91*), flow stretching (*11, 92*), and acoustic force spectroscopy (*93*). Despite these advances, a high-throughput, multimodal correlative single-molecule assay has remained out of reach, forcing researchers to maintain separate force, fluorescence, and correlative instruments. An integrated, scalable platform combining these modalities would resolve this fragmentation and define a new operating regime for single-molecule biophysics.

Several constraints guide the design of such a platform. For multiplexed single-molecule fluorescence, TIRF microscopy offers superior signal-to-noise over epifluorescence and supports low-nanomolar concentrations of fluorescently labeled biomolecules in solution (*3*). Among manipulation techniques, optical tweezers do not multiplex well, whereas camera-based, surface-tethered approaches are inherently scalable (*94*). Flow stretching offers experimental simplicity, but its micron-scale tethers limit spatial and temporal resolution. MT, by contrast, support both high throughput (*89*) and high spatiotemporal resolution (*4*), and integrate naturally with TIRF microscopy (*32, 36*). The combination of MT and TIRF therefore stands out as a particularly powerful route to high-throughput, multimodal correlative single-molecule measurements at the highest spatiotemporal resolution.

Here, we present such a platform, combining high-throughput magnetic tweezers with two-color TIRF microscopy to deliver force, torque, fluorescence, temperature-dependent and correlative measurements at single-molecule resolution (**Fig. 1**). We established objective-based TIRF with homogeneous illumination across an ultra-wide field of view that fully spans the sCMOS detector, enabling simultaneous monitoring of thousands of surface-immobilized fluorescent molecules and supporting accurate FRET measurements (**Fig. 2a**). We applied the assay to monitor the temperature-dependent FRET dynamics of a DNA hairpin between its folded and unfloded states with high temporal resolution (40 Hz image acquisition, **Fig. 2c**). Prism-based spectral imaging maximized the number of tethers imaged simultaneously on the MT and TIRF cameras, supporting correlative MT–TIRF measurements on up to ∼400 tethered magnetic beads in parallel (**Fig. 4**). We used this configuration to resolve the kinetics of the closed-to-open promoter complex transition during bacterial transcription initiation, illustrating the ability of the platform to interrogate protein–nucleic acid interactions across multiple single-molecule modalities (**Fig. 5**).

We anticipate that this platform will integrate high-throughput single-molecule assays into existing workflows that use next-generation sequencing to interrogate biomolecular reactions across sequence space (*6*) and cryo-EM to link structure and function. The now achievable throughput will open the door to large, multi-factor biomolecular complexes, such as mammalian systems, where low *in vitro* activity yield makes single-molecule analysis notoriously difficult to perform, extending the reach of these techniques to medically relevant systems. Parallel advances in high-throughput microfluidics, which enable increasingly parallelized experiments in extremely small volumes, place a complementary demand on scalable single-molecule detection (*92*). Expanding the multimodal imaging area will permit rapid interrogation of multiple conditions in a single experiment, and ongoing advances in large-area, high-speed, high-sensitivity sCMOS detectors should further extend throughput and spatiotemporal resolution, providing simultaneous access to fast and rare biomolecular events. Combined with automated acquisition and machine-learning data analysis, these advances should move scalable multimodal single-molecule biophysics from proof-of-principle into application-driven regimes such as drug discovery (*95*).

## Methods

### Optomechanics, optics, and electronics

The setup is a bespoke inverted microscope with two illuminations and imaging paths for both MT and TIRF channels, respectively (see **Fig. 1** and **Fig. S1a** for the schematics). In the following sections, each part is described separately.

### MT design

The magnetic field is generated by a pair of vertically aligned permanent magnets (5 mm cubes, W-05-N50-G, SuperMagnete, Switzerland) separated by a 1 mm gap which are located above the flow chamber. The vertical position and rotation of the magnets are actuated by the M-126-PD1 and C-150-PD motors controlled by C-863.11 servo drivers (Physik Instrumente). In **Fig. S1a,** the z-position of the microscope objective (Zeiss TIRF objective, oil immersion, 63×, NA=1.46) is precisely adjusted by a high-resolution piezo stage PIFOC (P-726.1CD, 100 µm travel range, controlled by the E-754.1CD controller, Physik Instrumente), while the sample position is coarsely adjusted by a mechanical z-stage (a few mm travel range with 10 µm resolution). The field of view is illuminated through the magnets gap by a collimated LED light source (Cree LED SGH, 460 nm) located above the magnets.

The magnetic and reference beads in the field of view are imaged onto a large-area sensor CMOS camera (Dalsa Falcon2 FA-80-12M1H, Stemmer Imaging) by the objective and a Zeiss tube lens (L3 in **Fig. S1a**, f = 165mm). All magnetic tweezers data were acquired at 58 Hz camera acquisition frequency. A dichroic mirror separates the MT illumination from the TIRF excitation (DM2 in **Fig. S1a**).

### TIRF design

#### TIRF excitation pathway

A BrixX laser box (Omicron Laserage Laserprodukte GmbH) provides TIRF excitation at 532 nm and 638 nm wavelengths with respective maximum output power of 1250 mW and 1200 mW. The lasers are coupled into a 50 µm step-index multimode fiber (MMF, FC/PC connector, and ceramic Ferrule). The output of the MMF is coupled to a cage optical system using a reflective collimator (RC) and further expanded with a beam expander consisting of a lens pair (BE in **Fig. S1a**), with respective focal lengths of-50 mm and 75 mm. The beam is then cropped by an adjustable iris to a diameter of

∼17 mm. Lens L1 (f = 150 mm) focuses the excitation lasers on the back focal plane (BFP) of the microscope objective after getting spectrally cleaned by an excitation filter (ExF in **Fig. S1a**). A translation stage enables the precise adjustment of the lateral position of the excitation beam away from the optical axis to enable both epifluorescence and TIRF excitation modes.

#### Speckle reduction unit

To eliminate the speckle pattern, a piezo element vibrating at a frequency of ∼1.3 kHz and a mechanical vibrator were employed (**Fig. S2a**). **Fig. S2b** describes how different propagation modes are affected by the bending of the MMF, resulting in mode scrambling when the fiber is vibrated. **Fig. S2c** shows the speckle contrast as a function of the piezo vibration frequency for different number of attachment points of the piezo (four in **Fig. S2a**), demonstrating a clear reduction in contrast at ∼1.3 kHz for the green laser. Speckle patterns for the 638 nm laser are intrinsically low and not altered by the vibrations. **Fig. S2de** show images of the speckle pattern for both laser wavelengths when applying the vibrator unit. The images are acquired using the assay described in **Fig. S2h**, where the output of the fiber is imaged by a lens pair forming an image of the fiber output onto a CMOS camera (Thorlabs, DCC1545M). The number on the corner for each beam intensity profile indicates the calculated speckle contrast. The characterization of the laser excitation homogeneity over the entire field of view in the TIRF setup is achieved by using TetraSpeck fluorescent beads, using either the 532 nm laser (30 W/cm^2^, 100 ms exposure time) or the 638 nm laser (30 W/cm^2^, 100 ms exposure time) with the vibration unit either off or on (**Fig. S2fg**).

#### TIRF Imaging pathway

The fluorescence emission from the sample is separated from the excitation source by a dichroic mirror (DM1, Chroma Tech. corp., **Fig. S1d**) and was further spectrally cleaned using an emission filter (**Fig. S1a**). A wedge prism spectrally decomposes the fluorescence-emitted photons to different angles (**Fig. S4a** and **Supplementary Information**), which steers the imaging optical path. A kinematic mirror placed in front of the camera steers the imaging optical path onto the sCMOS sensor. The image is formed on the sCMOS camera by a Zeiss tube lens (L2 in **Fig. S1a**, f = 165 mm).

### Controller unit

To control the vibrators and trigger the camera and the lasers, we used a microcontroller (Arduino Due®, 32-bit ARM core) as MCU that was programmed to synchronize both the illumination and acquisition for the TIRF and MT channels with sub-10 µs time resolution. The lasers in the TIRF channel can be triggered by user-defined patterns, including ALEX (*96*).

### DNA construct fabrication

#### Short duplex DNA assembly for single-molecule FRET experiments

The labeled 44 bp dsDNA constructs were assembled by annealing equimolar amounts of two complementary oligonucleotides (Biomers GmbH, sequence in **Supplementary Information**) in elution buffer (10 mM Tris-HCl, pH 8, 0.5 M NaCl) to the final concentration of 100 nM. The solution was heated up to 90°C for 1 min and then slowly cooled down to 4°C at a rate of 0.5 °C/min using a thermocycler (Bio-Rad, PTC Tempo).

#### DNA FRET Hairpin

A DNA hairpin for smFRET experiments was assembled according to Baltierra-Jasso et al. (*62*). In short, the DNA hairpin was assembled from oligos H2 and A2 (sequence in **Supplementary Information**). The oligos were synthesized and fluorescently labeled by biomers.net GmbH and annealed in an equimolar ratio in 20 mM Tris-HCl pH 8.0 and 50 mM NaCl at a final concentration of 10 µM. The sample was heated to 70°C for 5 min and subsequently cooled by 2°C/5 min to ensure accurate hybridization.

### 3.2 kbp dsRNA

To perform the intercalator benchmark experiment and evanescent field penetration depth measurement, we used a 3.2 kbp coilable double-stranded RNA construct, the fabrication is described in detail in Ref. (*71*).

### 21 kbp DNA

To perform the magnetic tweezers force calibration, we used a 20,666 bp construct, whose fabrication was previously described in detail in Ref. (*45*) and the sequence is provided in Ref. (*97*).

### DNA construct to monitor RPO dynamics

To perform the RPO experiments, a linear ∼1.4 kbp long double-stranded DNA construct encoding the lacCONS+2 promoter sequence was obtained by digestion and ligation as described previously (*71, 97*). Shortly, digoxigenin-or biotin-labeled DNA handles (394 and 415 bp, respectively) were generated by Taq DNA polymerase PCR (NEB) with modified dUTPs, either biotin-16-dUTP or digoxigenin-11-dUTP (Jena Biosciences GmbH). The DNA fragment that encodes the lacCONS+2 promoter was amplified by PCR using the Phusion High-Fidelity DNA polymerase (NEB). DNA fragments were digested for 2 hours at 37 °C. The ligation of the digested DNA handles and stem was performed using T4 DNA ligase (NEB) at 16°C overnight. The sequences of the primers and construct are provided in **Supplementary Information**.

### Expression, purification, and labeling of transcription initiation components

*Escherichia coli* RNA polymerase was purified by using sequentially nickel affinity, heparin affinity and anion exchange chromatography steps as previously described (*98*).The expression plasmid for *Escherichia coli σ*^70^ was constructed by amplifying *rpoD* by PCR from the genomic DNA isolated from *E. coli* K-12 and using Gibson assembly to insert the product into multiple cloning site of pET28b (*99*). The construct, named as pRP041, includes His tag followed by a cut site for TEV protease at the N-terminal front of *rpoD*. To allow site-specific attachment of a fluorophore to Cys132 in *σ*^70^, codons encoding two other Cys residues (at amino acid positions 291 and 295) in wild type *rpoD* were substituted to encode serines; the resulting plasmid was named as pAZ003. The region containing the *rpoD* reading frame, T7 promoter and T7 transcription terminator was verified by Sanger sequencing in final plasmids (Eurofins Genomics). The plasmids were maintained and purified from *E. coli* XL1-blue cells (Stratagene), which were cultivated in LB medium supplemented with 30 µg/ml kanamycin.

To produce the single cysteine *σ*^70^ (C291S, C295S), pAZ003 was transformed into *E. coli* C41(DE3) cells (*100*). For protein production the culture was grown in two 2 liters Erlenmeyer flasks each containing 0.5 l of LB medium supplemented with 30 µg/ml kanamycin. The culture was grown at 30 °C in an orbital shaker set to 250 rpm (Multitron, Infors HT) until absorbance at 600 nm reached 0.6, at which point the culture temperature was dropped to 16 °C and 0.4 mM IPTG was added to induce *σ*^70^production. The cultivation was continued for about 16 h before the cells were harvested by centrifugation at 3000 *g,* cells were frozen and stored at-80 °C. The cell pellet was resuspended in lysis buffer (30 mM Tris-HCl, pH 7.0, 500 mM NaCl, 5% glycerol, 1 mM Tris [2-carboxyethyl] phosphine [TCEP]) supplemented with 0.2% Tween-20, 10 mM imidazole, 5 mg lysozyme and protease inhibitors [1 tablet of Pierce™ Protease Inhibitor Mini Tablets, EDTA-free; Thermo Fisher Scientific; cat.no. A32955]. Cells were lysed by sonication, and the lysate was clarified by centrifugation at 70 000 *g* for 1 h. The supernatant was passed through a PD10 gravity column (Cytiva) packed with about 1 ml bed of PureCube 100 Ni-NTA agarose (Cube Biotech). The matrix was then washed with lysis buffer supplemented with 20 mM imidazole to remove weakly bound proteins, followed by the elution of *σ*^70^ using lysis buffer supplemented with 500 mM imidazole. The eluate was diluted with Buffer A (20 mM MES, pH 6.5, 5% glycerol, 1 mM Tris [2-carboxyethyl] phosphine [TCEP], 0.1 mM EDTA) to a final NaCl concentration of 80 mM. Sample was applied to a 5 ml HiTrap Heparin HP (Cytiva) column. The heparin column was developed at 1 ml/min flow rate using ÄktaPrime Plus system (Cytiva) and NaCl gradient in Buffer B (20 mM MES, pH 6.5, 1500 mM NaCl, 5% glycerol, 1 mM Tris (2-carboxyethyl) phosphine [TCEP], 0.1 mM EDTA) to elute *σ*^70^, which occurred at about 350 mM NaCl. As the final purification step, the *σ*^70^ containing fractions from the heparin step were combined, diluted with Buffer A to a final NaCl concentration of 80 mM. Sample was applied to a 6 ml RESOURCE Q anion exchange column (Cytiva). The anion exchange column was developed as above but using Äkta Purifier system (Cytiva); *σ*^70^ eluted in NaCl gradient in Buffer B at about 320-350 mM NaCl. The final sample was concentrated with 3 kDa Ultra Centrifugal Filter (Amicon), to a final concentration of 133 µM, flash-frozen in liquid nitrogen and stored at-80°C.

For labeling *σ*^70^ the sample buffer (20 mM MES, pH 6.5, 350 mM NaCl, 5% glycerol, 1 mM Tris (2-carboxyethyl) phosphine [TCEP], 0.1 mM EDTA) was replaced using 0.5 ml Zeba™ Spin Desalting Column (7kDa cutoff, Thermo Fisher Scientific) with Dilution buffer (20 mM Tris-HCl, pH 7.5, 500 mM NaCl, and 10% glycerol). Maleimide conjugated ATTO550 dye (ATTO-TEC) was dissolved dimethylformamide at concentration 10 mg/ml. 120 µM *σ*^70^ and 1300 µM ATTO550 dye were then mixed in 50 µL total volume (in Dilution buffer) and incubated for 2 h at 20°C with gentle stirring every half an hour. The reaction was quenched by adding 10 mM dithiothreitol to the mixture. The free dye was then removed by pushing the quenched reaction mixture through a 0.5 ml Zeba™ Spin Desalting Column (7kDa cutoff, Thermo Fisher Scientific). The residual free dye was further removed by dialyzing the *σ*^70^ sample overnight at 4°C against 250 ml of Dilution buffer supplemented with 2 mM dithiothreitol. The final *σ*^70^ sample was aliquoted, flash-frozen in liquid nitrogen and stored at-80°C. The labeling efficiency of *σ*^70^ was evaluated by estimating total protein and dye concentrations from light absorbance at 280 and 554 nm, respectively. The used molar absorption coefficients (*ε*) were 42860 M^-1^ cm^-1^ and 120 000 M^-1^ cm^-1^ for *σ*^70^ and ATTO550, respectively. For protein concentration estimation, the absorbance of ATTO550 at 280 nm, obtained by multiplying the absorbance of ATTO550 at 550 nm by 0.1, was subtracted from total absorbance at 280 nm. The success of *σ*^70^ labeling was further verified by SDS-PAGE in conjunction with the gel scanning with Sapphire Biomolecular Imager (Azure Biosystems) using the fluorescence mode. Data indicated 99% labeling efficiency of *σ*^70^.

### Flow chamber assembly and preparation

#### Flow chamber assembly for experiments including magnetic tweezers measurements

The flow chamber is made of two coverslips (top and bottom) sandwiching a spacer made of a 2-layer thick Parafilm (Sigma Aldrich) with a carved channel. The bottom coverslip is low-fluorescence (Epredia, #1.5, 24×60 mm) and the top coverslip is #1 thickness (24×60 mm, #631-1339, Menzel Glaser, VWR) that has two holes drilled using a tabletop sandblaster (Vaniman) (*101*). Before assembly, the top and bottom coverslips were washed by sonication in a solution of Hellmanex III (Sigma-Aldrich) diluted in deionized water (∼2% v/v) and subsequently rinsed thoroughly under a stream of deionized water and dried in an oven at 80°C and stored in a 50 ml Falcon tube. Coverslips were activated and made hydrophilic through thorough surface treatment using a Corona SB discharger (Electro-Technic Products). The bottom coverslip was subsequently treated for surface functionalization as described in the following paragraph depending on the experiment type and needs. The flow chamber is subsequently sealed by melting the Parafilm on a hot plate at ∼100 °C.

#### Reference beads surface attachment for experiments including magnetic tweezers

A diluted stock of 2 µm polystyrene reference beads was prepared by diluting 1 µl stock (Sigma, Cat # 78452) 1000-fold in deionized water and rinsing them by repeating the following procedure three times: vortex, centrifugation at ∼13000 rpm for ∼1 min, removal of the supernatant and resuspension in the same volume of deionized water. The washed reference beads were finally resuspended in 1000 µl of deionized water and stored at 4°C for further use. For MT experiments, preceding the assembly of the flow cell, ∼3 µl of the reference bead solution was spread with the side of a pipet tip on the top surface of the bottom coverslip of the flow cell, and subsequently heated to ∼100°C for ∼5 min to melt the reference beads on the coverslip surface.

#### Surface preparation for experiments including magnetic tweezers

We used two different types of surface passivation to perform our experiments that include magnetic tweezers; using BSA or lipid bilayer, which are described below.

#### BSA passivated flow chamber

First, 50 µl of full-length anti-digoxigenin (0.5 mg/ml in Phosphate buffered saline (PBS), Sigma Aldrich, preheated at 37 °C for ≥20 min) was incubated in the flow cell for 30 min. The excess was subsequently flushed away with 1 ml of PBS buffer containing 700 mM NaCl followed by 10 min incubation, and a rinsing step with 1 ml PBS buffer. Then the flow chamber was incubated for ∼5 minutes with a mixture of 5 mg/ml bovine serum albumin (NEB) diluted in TE150 buffer (10 mM Tris, 1mM EDTA, 150 mM NaCl and 2 mM sodium azide). The excess proteins were subsequently flushed out with ∼1 ml of TE150.

This method is used for the force calibration, rotation calibration and the benchmark experiment using intercalators.

#### SUV solution preparation for lipid bilayer-functionalized flow chamber

The small unilamellar vesicles (SUVs) were prepared based on the previously published literature (*102*). We first mixed DOPC (1,2-dioleoyl-sn-glycero-3-phosphocholine) and PEG-PE (1,2-dioleoyl-sn-glycero-3-phosphoethanolamineN-[methoxy(polyethyleneglycol)-550]) (850375C, 880530C, respectively, Avanti Polar Lipids) dissolved in chloroform in 95:5 molar fraction. After mixing, the solution was dried under nitrogen flow, followed by drying in a vacuum desiccator for 1 h. The dry lipid film was then resuspended at 4 mg/ml in the storage buffer (10 mM HEPES, pH 7.8, 150 mM NaCl, 2 mM EDTA, 2 mM sodium azide) by vortexing. Then the suspension was incubated for 30 min at room temperature, with occasional vortexing and subsequently flash frozen-thawed 5 times using liquid nitrogen and a heating block. SUV stock solution was aliquoted and stored at-20°C, maintaining their functionality for several weeks.

#### Lipid bilayer passivated flow chamber

As described in detail in ref. (*97, 101*). Following assembly, the flow cell was rinsed with 1 ml 1x PBS. 50 µl of full-length anti-digoxigenin (0.5 mg/ml in PBS, Sigma Aldrich, preheated at 37 °C for ≥20 min) was incubated in the flow cell for 30 min. The excess was subsequently flushed away with 1 ml of 1x PBS buffer containing 700 mM NaCl followed by 10 min incubation, and a rinsing step with 1 ml PBS. The buffer in the flow cell was exchanged with 1 ml vesicle dilution buffer (10 mM HEPES pH 7.4, 150 mM NaCl, 2 mM EDTA, 2 mM sodium azide and 2 mM CaCl2) to prepare the lipid bilayer assembly. The lipid bilayer was formed by flushing 1.2 ml of the SUV solution (described below) at 50 µg/ml in the vesicle dilution buffer (Sigma Aldrich), slowly through the flow cell at 0.1 ml/min for a total duration of ∼12 min. The flow channel was then washed with 1 ml PBS to remove any excess SUVs. To finalize the surface passivation, BSA (NEB) at 1 mg/ml in PBS was incubated for 30 min and subsequently flushed away excess BSA with 1 ml PBS.

This method is used for evanescent field penetration depth measurement and RPO dynamics experiment.

#### Flow chamber assembly and preparation for TIRF-only experiments

Glass cover slides (Epredia, #1.5, 24×60 mm, Ted Pella, Inc.) and reusable slide gaskets (GBL103280, Merck) were cleaned via sonication in Hellmanex III (Sigma-Aldrich) and deionized water. The glass cover slides were plasma treated before placing a gasket on the surface. The reaction chamber was functionalized with 1 mg/ml recombinant albumin (NEB), and 0.02 µg/ml Biotin-BSA (Sigma-Aldrich) in TE150 (10 mM Tris-HCl pH 8.0, 1 mM EDTA, and 150 mM NaCl) for 5 min. After rinsing the gasket with TE150. The gasket was passivated with 4 mg/ml recombinant albumin, and 20 µg/ml Neutravidin (Thermo Fisher Scientific) in TE150. After 5 min incubation the surface was again thoroughly rinsed with TE150. The surface was subsequently rinsed with 20 mM Tris-HCl and 15 mM NaCl.

This method is used for the short duplex DNA experiment, and for the DNA FRET hairpin experiment.

#### TetraSpeck™ microspheres experiments

The top surface of the bottom coverslip of the flow chamber was coated with Nitrocellulose (Sigma Aldrich) by spreading a 5 µl droplet of nitrocellulose dissolved in pentylacetate (0.1% w/v; Sigma Aldrich) over the coverslip. After evaporation of the pentylacetate, the nitrocellulose-coated surface was used as the top surface of the bottom coverslip of the flow chamber, in which double layer of Parafilm has been sandwiched between the coverslips on a 100°C hot plate. The flow chamber was then incubated with 100x diluted (in deionized water) TetraSpeck™ microspheres (Thermo Fisher Scientific, ∼10^8^ particles/ml, fluorescent orange, #L5530) for 10 minutes and excess beads were subsequently flushed away with ∼1 ml of TE150 buffer.

### Imaging buffers

#### Short duplex DNA, benchmark using intercalator, and RPo dynamics experiments

Imaging was performed with the sample in an imaging buffer (50 mM HEPES pH 7.9, 100 mM potassium glutamate, 10 mM DTT, and 10 mM MgCl2) containing 1 mM cyclooctatetraene (COT) and 1 mM Trolox as a triplet-state quencher, and Gloxy (10 mg/ml glucose, 1 mg/ml glucose oxidase (Sigma-Aldrich), 0.4 µg/ml catalase, 0.1 mM Tris and 0.5 mM NaCl) as oxygen scavenger, as indicated. The Trolox was prepared by dissolving 2.5 mg Trolox (Sigma-Aldrich) in 1 ml of Tris (pH ∼8.0), then placed under sunlight for 1 or 2 days to see a color change, then made aliquots, and stored at-20°C. The 100 mM COT was prepared by dissolving stock solution (Sigma-Aldrich) in DMSO and stored at-20 °C.

#### DNA FRET hairpin experiment

Labeled hairpin experiments were performed in an oxygen scavenger buffer containing 20 mM Tris-HCl pH 8.0, 15 mM NaCl, 0.2 mg/ml recombinant albumin, 5 U/ml Bilirubin oxidase (Merck), 10 mM sodium ascorbate (Merck), and 2 mM Trolox (*103*). The Trolox solution was prepared as described above.

#### Other experiments

Other experiments including force calibration, measurements on TetraSpeck, rotation calibration and measurements on evanescent field penetration depth are imaged using TE150 buffer.

### Data recording

The MT and TIRF images were processed in real-time and simultaneously using a custom Python interface. This interface includes trackers in pytorch, CUDA, and C++ to enable real-time processing of both the magnetic beads 3D positions and fluorescence spots intensity and position using a graphics processing unit (GPU) card to enable real-time analysis. The magnetic beads 3D positions are determined using the GPU-accelerated localization algorithms based on the quadrant interpolation (QI) previously described in Ref. (*14*).

For TIRF spot tracking, we developed two separate methods utilizing a custom-made point spread function (PSF) template, in which the PSF shape and inter-spot distance is estimated by averaging over spot pairs for low-FRET oligo sample when imaged in ALEX mode (**Fig. S4c**). The first method integrates pixel intensities within the predefined spot boundaries defined by the PSF template such that ∼95% of the total emission of each dye is captured, and the background defined by averaging over the pixels outside the template, offering a computationally efficient solution when spot positions are fixed. Spot localization is performed for the first step detection by finding local maxima via cross-correlation with the template in the frequency domain. This approach is suited for high-throughput TIRF measurements involving thousands of single-molecule fluorescence spots. This approach is used for calibration measurements of low-and high-FRET duplex oligos and for the measurements on the hairpin kinetics. Alternatively, in applications requiring higher precision in localization, we employed maximum likelihood estimation (MLE) fitting. In this case, a Levenberg–Marquardt optimization algorithm is used to determine the parameters (spot position, intensity and background) that maximize the log-likelihood function under the assumption of Poisson-distributed photon counts. This latter approach is used for correlative MT–TIRF measurements where the number of TIRF spots is limited by the number of tethers.

### Sample incubation and data acquisition conditions

#### Short duplex DNA experiments

For the short duplex DNA experiment, the sample was applied to the gasket at ∼100 pM and incubated for ∼10 sec until an acceptable coverage is observable on the TIRF channel and subsequently rinsed with 20 mM Tris-HCl.

The laser intensities were measured above the objective in epifluorescence illumination and set to ∼100 W/cm^2^ for both lasers. The sCMOS camera exposure time was set to 1000 ms. The laser illumination timing was set to cover the full camera acquisition period in the alternating laser excitation (ALEX) scheme.

#### Temperature-dependent DNA hairpin dynamics experiments

For the DNA FRET hairpin experiment, the sample was applied to the gasket at 100 pM and incubated for ∼5 sec and subsequently rinsed with 20 mM Tris-HCl, 15 mM NaCl, and 0.2 mg/ml recombinant albumin.

Images were recorded at 40 Hz and 25 ms exposure time. The donor dye was excited with 200 W/cm^2^ at 532 nm. Dynamics of the hairpin opening and closing were recorded for approximately 3000 frames at 24 °C to 30 °C.

#### High-throughput correlative benchmark experiment using dsRNA and SYBR^TM^ Gold

To make the tethers, 10 µl of the streptavidin-coated superparamagnetic beads (NEB, S1420S, 1 µm) were washed three times in TE150 buffer and diluted to 50 µl of TE150 buffer mixed with ∼10 pM of the ∼3.2 kbp coilable dsRNA construct (*71*) and incubated for a few minutes. The dsRNA-tethered magnetic beads were then flushed into the flow cell and incubated for ∼5 min to ensure attachment of the digoxigenin-labeled RNA handle to the anti-digoxigenin adsorbed on the flow cell surface. Finally, the excess magnetic beads were removed by flushing copious amounts of TE150 buffer with occasional gentle tapping on the exit tube connecting the flow cell to the withdrawing pump. 500 nM SYBR™ Gold mixed in the imaging buffer (Gloxy, as described earlier) was added with the rate of 1 µl/min while the sample was imaged by MT (58 Hz) and TIRF (1 Hz, 100 ms exposure time) channels using 50 W/cm^2^ 532nm laser illumination.

#### RNA polymerase-promoter open complex (RPO) experiment

To make the tethers, 10 µl of the streptavidin-coated superparamagnetic beads (NEB, S1420S, 1 µm diameter) were washed 3× in PBS and diluted to 50 µl of PBS containing ∼30 pM of DNA construct containing the lacCONS+2 promoter for Escherichia coli RNA polymerase (see DNA construct fabrication section and ref. (*79*)), and incubated for a few minutes. The dsDNA-tethered magnetic beads were then flushed into the flow cell and incubated for ∼5 min to ensure attachment of the digoxigenin-labeled DNA handle to the anti-digoxigenin adsorbed on the flow cell surface. Finally, the excess magnetic beads were removed by flushing copious amounts of PBS with occasional gentle tapping on the exit tube connecting the flow cell to the withdrawing pump.

Holoenzymes were assembled by incubating 0.5 µM RNA polymerase with 1 µM labeled *σ*^70^ (at the ratio of 1:2 core:sigma factor) in 20 mM Tris-HCl pH 7.9, 150 mM NaCl, 0.1 mM EDTA, 50% (V/V) Glycerol, and 0.1 mM DTT, and incubated at 30 °C for 30 min as described in Ref. (*79*). The resulting holo-complex was stored at – 20 °C.

The extension and coilability of the DNA molecule were evaluated by stretching the DNA from 0to 4 pN and rotating the magnetic bead first in negative and then in positive directions (±12 turns) while keeping the force at 4 pN. For a coilable single DNA tether molecule, the extension should remain constant in the negative turn region but decrease in the positive turn region (*15, 97*).

The flow chamber temperature was set to 34 °C using the objective-based temperature control we previously described (*47*). For the control measurement, we used imaging buffer containing Gloxy and 0.5 mg/ml BSA, and 150 mM KGlu without the holoenzyme in the flow chamber first, and subsequently with 5 nM of holoenzyme added to the aforementioned reaction mixture and flushed at 0.1 ml/min while the tethers were stretched by 4 pN and rotated +3 turns to ensure the beads are clamped to prevent bead rotation and open complex formation during the flush. Force was then decreased to ∼0.3 pN and the magnets rotated to-4 turns position to start the experiment. The MT data were recorded at 58 Hz acquisition frequency. In parallel, the fluorescence images were acquired, with an exposure time of 100 ms (3.3 frames per second) and 100 W/cm^2^ on the field of view of the 532 nm laser. The CMOS and sCMOS cameras, the laser, and LED illuminations were synchronized using the Arduino controller unit.

## Acknowledgments

DD was supported by the Interdisciplinary Center for Clinical Research (IZKF) at the University Hospital of the University of Erlangen-Nuremberg, and the German Research Foundation grant DFG-DU-1872/3-1, DFG-DU-1872/5-1, and BaSyC – Building a Synthetic Cell” Gravitation grant (024.003.019) of the Netherlands Ministry of Education, Culture and Science (OCW) and the Netherlands Organization for Scientific Research (NWO) and NWO-M Open Competition Domain Science grant OCENW.M.21.184 and OCENW.M.23.350. AMM was supported by Sigrid Jusélius foundation. We thank current and former members of the Dulin lab (Subhas C. Bera, Flavia S. Papini, Luca Buccolieri, Pim America, Paula Rivas, Misha Klein) for assisting with the project. We thank Chirlmin Joo’s lab for assistance with the surface chemistry.

## Contributions

DD designed and led the research. DD, YW and MSF designed the instrument. YW and MSF built the instrument, designed and assembled the electronics control, with the assistance from TB. JC and MSF contributed custom software. DD and MSF designed the experiments. MSF, SQ, and AR performed the experiments and MSF and SQ analyzed them. QS and SQ contributed DNA and RNA constructs. AZ, RKP and AMM contributed the fluorescently labeled *σ*70 and the core RNAP. DD and MSF wrote the manuscript. All the authors edited the manuscript.

## Data availability

The data of this study will be made available upon acceptance of the manuscript though a DOI link.

## Supplementary Data

Supplementary Data are available online.

## References

1. B. P. English, W. Min, A. M. van Oijen, K. T. Lee, G. Luo, H. Sun, B. J. Cherayil, S. C. Kou, X. S. Xie, Ever-fluctuating single enzyme molecules: Michaelis-Menten equation revisited. Nature chemical biology 2, 87–94 (2006).

2. S. N. Xie, Single-molecule approach to enzymology. Single Mol 2, 229–236 (2001).

3. D. Dulin, J. Lipfert, M. C. Moolman, N. H. Dekker, Studying genomic processes at the single-molecule level: introducing the tools and applications. Nature reviews. Genetics 14, 9–22 (2013).

4. D. Dulin, B. A. Berghuis, M. Depken, N. H. Dekker, Untangling reaction pathways through modern approaches to high-throughput single-molecule force-spectroscopy experiments. Curr Opin Struct Biol 34, 116–122 (2015).

5. F. R. Hill, E. Monachino, A. M. van Oijen, The more the merrier: high-throughput single-molecule techniques. Biochem Soc Trans 45, 759–769 (2017).

6. A. N. Kapanidis, L. Muras, K. Sreenivasa, J. P. Hazra, J. van Noort, C. Joo, S. Deindl, From sequence to function: Bridging single-molecule kinetics and molecular diversity. *Science (New York*, N.Y 391, 458–465 (2026).

7. J. Reeks, P. Mahajan, M. Clark, S. R. Cowan, E. Di Daniel, C. P. Earl, S. Fisher, R. S. Holvey, S. M. Jackson, E. Lloyd-Evans, C. M. Morgillo, P. N. Mortenson, M. O’Reilly, C. J. Richardson, P. Schöpf, D. M. Tams, H. Waller-Evans, S. E. Ward, S. Whibley, P. A. Williams, C. N. Johnson, High throughput cryo-EM provides structural understanding for modulators of the lysosomal ion channel TRPML1. Structure 33, 1374–1385.e1377 (2025).

8. Y. Li, J. N. Cash, J. J. G. Tesmer, M. A. Cianfrocco, High-Throughput Cryo-EM Enabled by User-Free Preprocessing Routines. Structure 28, 858–869.e853 (2020).

9. I. Heller, G. Sitters, O. D. Broekmans, G. Farge, C. Menges, W. Wende, S. W. Hell, E. J. Peterman, G. J. Wuite, STED nanoscopy combined with optical tweezers reveals protein dynamics on densely covered DNA. Nature methods 10, 910–916 (2013).

10. M. J. Comstock, T. Ha, Y. R. Chemla, Ultrahigh-resolution optical trap with single-fluorophore sensitivity. Nature methods 8, 335–340 (2011).

11. K. E. Duderstadt, H. J. Geertsema, S. A. Stratmann, C. M. Punter, A. W. Kulczyk, C. C. Richardson, A. M. van Oijen, Simultaneous Real-Time Imaging of Leading and Lagging Strand Synthesis Reveals the Coordination Dynamics of Single Replisomes. Mol Cell 64, 1035–1047 (2016).

12. D. Dulin, “An Introduction to Magnetic Tweezers” in Single Molecule Analysis: Methods and Protocols, I. Heller, D. Dulin, E. J. G. Peterman, Eds. (Springer US, New York, NY, 2024), pp. 375–401.

13. D. Dulin, T. J. Cui, J. Cnossen, M. W. Docter, J. Lipfert, N. H. Dekker, High Spatiotemporal-Resolution Magnetic Tweezers: Calibration and Applications for DNA Dynamics. Biophysical journal 109, 2113–2125 (2015).

14. J. P. Cnossen, D. Dulin, N. H. Dekker, An optimized software framework for real-time, high-throughput tracking of spherical beads. The Review of scientific instruments 85, 103712 (2014).

15. T. R. Strick, J. F. Allemand, D. Bensimon, A. Bensimon, V. Croquette, The elasticity of a single supercoiled DNA molecule. *Science (New York*, N.Y 271, 1835–1837 (1996).

16. A. Lof, P. U. Walker, S. M. Sedlak, S. Gruber, T. Obser, M. A. Brehm, M. Benoit, J. Lipfert, Multiplexed protein force spectroscopy reveals equilibrium protein folding dynamics and the low-force response of von Willebrand factor. Proceedings of the National Academy of Sciences of the United States of America 116, 18798–18807 (2019).

17. S. Quack, S. Maity, P. P. B. America, M. Klein, A. Herrero del Valle, R. Singh, J. D. Joiner, Z. M. Rashid, Q. Smitskamp, P. Rivas, Flavia S. Papini, C. P. Broedersz, W. H. Roos, Y. Modis, D. Dulin, MDA5 generates compact ribonucleoprotein complexes via ATP-dependent double-stranded RNA unwinding. Nucleic acids research 54, gkag274 (2026).

18. P. P. B. America, S. C. Bera, A. Das, T. K. Anderson, J. C. Marecki, F. S. Papini, J. J. Arnold, R. N. Kirchdoerfer, C. E. Cameron, K. D. Raney, M. Depken, D. Dulin, RNA virus polymerase-helicase coupling enables rapid elongation through duplex RNA. Cell Rep 45, 117273 (2026).

19. M. Manosas, X. G. Xi, D. Bensimon, V. Croquette, Active and passive mechanisms of helicases. Nucleic acids research 38, 5518–5526 (2010).

20. T. R. Strick, V. Croquette, D. Bensimon, Single-molecule analysis of DNA uncoiling by a type II topoisomerase. Nature 404, 901–904 (2000).

21. A. Revyakin, C. Liu, R. H. Ebright, T. R. Strick, Abortive initiation and productive initiation by RNA polymerase involve DNA scrunching. *Science (New York*, N.Y 314, 1139–1143 (2006).

22. D. A. Koster, K. Palle, E. S. Bot, M. A. Bjornsti, N. H. Dekker, Antitumour drugs impede DNA uncoiling by topoisomerase I. Nature 448, 213–217 (2007).

23. D. Dulin, J. J. Arnold, T. van Laar, H. S. Oh, C. Lee, A. L. Perkins, D. A. Harki, M. Depken, C. E. Cameron, N. H. Dekker, Signatures of Nucleotide Analog Incorporation by an RNA-Dependent RNA Polymerase Revealed Using High-Throughput Magnetic Tweezers. Cell Rep 21, 1063–1076 (2017).

24. M. Seifert, S. C. Bera, P. van Nies, R. N. Kirchdoerfer, A. Shannon, T.-T.-N. Le, X. Meng, H. Xia, J. M. Wood, L. D. Harris, F. S. Papini, J. J. Arnold, S. Almo, T. L. Grove, P.-Y. Shi, Y. Xiang, B. Canard, M. Depken, C. E. Cameron, D. Dulin, Inhibition of SARS-CoV-2 polymerase by nucleotide analogs from a single-molecule perspective. eLife 10, e70968 (2021).

25. L. Colson, Y. Kwon, S. Nam, A. Bhandari, N. M. Maya, Y. Lu, Y. Cho, Trends in Single-Molecule Total Internal Reflection Fluorescence Imaging and Their Biological Applications with Lab-on-a-Chip Technology. Sensors-Basel 23, 10.3390/s23187691 (2023).

26. C. Joo, H. Balci, Y. Ishitsuka, C. Buranachai, T. Ha, Advances in single-molecule fluorescence methods for molecular biology. Annual review of biochemistry 77, 51–76 (2008).

27. M. F. Juette, D. S. Terry, M. R. Wasserman, R. B. Altman, Z. Zhou, H. Zhao, S. C. Blanchard, Single-molecule imaging of non-equilibrium molecular ensembles on the millisecond timescale. Nature methods 13, 341–344 (2016).

28. M. F. Juette, D. S. Terry, M. R. Wasserman, Z. Zhou, R. B. Altman, Q. Zheng, S. C. Blanchard, The bright future of single-molecule fluorescence imaging. Current opinion in chemical biology 20, 103–111 (2014).

29. J. D. Larson, M. L. Rodgers, A. A. Hoskins, Visualizing cellular machines with colocalization single molecule microscopy. Chem Soc Rev 43, 1189–1200 (2014).

30. E. Lerner, T. Cordes, A. Ingargiola, Y. Alhadid, S. Chung, X. Michalet, S. Weiss, Toward dynamic structural biology: Two decades of single-molecule Forster resonance energy transfer. *Science (New York*, N.Y 359, (2018).

31. E. Lerner, A. Barth, J. Hendrix, B. Ambrose, V. Birkedal, S. C. Blanchard, R. Borner, H. Sung Chung, T. Cordes, T. D. Craggs, A. A. Deniz, J. Diao, J. Fei, R. L. Gonzalez, I. V. Gopich, T. Ha, C. A. Hanke, G. Haran, N. S. Hatzakis, S. Hohng, S. C. Hong, T. Hugel, A. Ingargiola, C. Joo, A. N. Kapanidis, H. D. Kim, T. Laurence, N. K. Lee, T. H. Lee, E. A. Lemke, E. Margeat, J. Michaelis, X. Michalet, S. Myong, D. Nettels, T. O. Peulen, E. Ploetz, Y. Razvag, N. C. Robb, B. Schuler, H. Soleimaninejad, C. Tang, R. Vafabakhsh, D. C. Lamb, C. A. Seidel, S. Weiss, FRET-based dynamic structural biology: Challenges, perspectives and an appeal for open-science practices. eLife 10, (2021).

32. E. T. Graves, C. Duboc, J. Fan, F. Stransky, M. Leroux-Coyau, T. R. Strick, A dynamic DNA-repair complex observed by correlative single-molecule nanomanipulation and fluorescence. Nature structural & molecular biology 22, 452–457 (2015).

33. F. E. Kemmerich, M. Swoboda, D. J. Kauert, M. S. Grieb, S. Hahn, F. W. Schwarz, R. Seidel, M. Schlierf, Simultaneous Single-Molecule Force and Fluorescence Sampling of DNA Nanostructure Conformations Using Magnetic Tweezers. Nano letters 16, 381–386 (2016).

34. Y. Seol, K. C. Neuman, Combined Magnetic Tweezers and Micro-mirror Total Internal Reflection Fluorescence Microscope for Single-Molecule Manipulation and Visualization. Methods Mol Biol 1665, 297–316 (2018).

35. M. Lee, S. H. Kim, S.-C. Hong, Minute negative superhelicity is sufficient to induce the B-Z transition in the presence of low tension. Proceedings of the National Academy of Sciences 107, 4985–4990 (2010).

36. P. Aldag, M. Rutkauskas, J. Madariaga-Marcos, I. Songailiene, T. Sinkunas, F. Kemmerich, D. Kauert, V. Siksnys, R. Seidel, Dynamic interplay between target search and recognition for a Type I CRISPR-Cas system. Nature communications 14, 3654 (2023).

37. J. Madariaga-Marcos, S. Hormeño, C. L. Pastrana, G. L. M. Fisher, M. S. Dillingham, F. Moreno-Herrero, Force determination in lateral magnetic tweezers combined with TIRF microscopy. Nanoscale 10, 4579–4590 (2018).

38. M. T. van Loenhout, M. V. de Grunt, C. Dekker, Dynamics of DNA supercoils. *Science (New York*, N.Y 338, 94–97 (2012).

39. Q. Guo, Y. He, H. P. Lu, Interrogating the activities of conformational deformed enzyme by single-molecule fluorescence-magnetic tweezers microscopy. Proceedings of the National Academy of Sciences 112, 13904–13909 (2015).

40. X. Long, J. W. Parks, C. R. Bagshaw, M. D. Stone, Mechanical unfolding of human telomere G-quadruplex DNA probed by integrated fluorescence and magnetic tweezers spectroscopy. Nucleic acids research 41, 2746–2755 (2013).

41. C. Lu, M. Talebi Khoshmehr, M. S. Feiz, M. M. Dashtabi, D. Dulin, B. Imran Akca, Entropy-based super-resolution imaging in waveguide-based TIRF microscopy&#x2014;an experimental and numerical study. Opt Express 33, 29295–29307 (2025).

42. R. Diekmann, Ø. I. Helle, C. I. Øie, P. McCourt, T. R. Huser, M. Schüttpelz, B. S. Ahluwalia, Chip-based wide field-of-view nanoscopy. Nature Photonics 11, 322–328 (2017).

43. B. Hellenkamp, S. Schmid, O. Doroshenko, O. Opanasyuk, R. Kühnemuth, S. Rezaei Adariani, B. Ambrose, M. Aznauryan, A. Barth, V. Birkedal, M. E. Bowen, H. Chen, T. Cordes, T. Eilert, C. Fijen, C. Gebhardt, M. Götz, G. Gouridis, E. Gratton, T. Ha, P. Hao, C. A. Hanke, A. Hartmann, J. Hendrix, L. L. Hildebrandt, V. Hirschfeld, J. Hohlbein, B. Hua, C. G. Hübner, E. Kallis, A. N. Kapanidis, J.-Y. Kim, G. Krainer, D. C. Lamb, N. K. Lee, E. A. Lemke, B. Levesque, M. Levitus, J. J. McCann, N. Naredi-Rainer, D. Nettels, T. Ngo, R. Qiu, N. C. Robb, C. Röcker, H. Sanabria, M. Schlierf, T. Schröder, B. Schuler, H. Seidel, L. Streit, J. Thurn, P. Tinnefeld, S. Tyagi, N. Vandenberk, A. M. Vera, K. R. Weninger, B. Wünsch, I. S. Yanez-Orozco, J. Michaelis, C. A. M. Seidel, T. D. Craggs, T. Hugel, Precision and accuracy of single-molecule FRET measurements—a multi-laboratory benchmark study. Nature methods 15, 669–676 (2018).

44. S. Quack, D. Dulin, “Surface Functionalization, Nucleic Acid Tether Characterization, and Force Calibration for a Magnetic Tweezers Assay” in Single Molecule Analysis: Methods and Protocols, I. Heller, D. Dulin, E. J. G. Peterman, Eds. (Springer US, New York, NY, 2024), pp. 403–420.

45. E. Ostrofet, F. S. Papini, D. Dulin, Correction-free force calibration for magnetic tweezers experiments. Sci Rep 8, 15920 (2018).

46. S. C. Bera, M. Seifert, R. N. Kirchdoerfer, P. van Nies, Y. Wubulikasimu, S. Quack, F. S. Papini, J. J. Arnold, B. Canard, C. E. Cameron, M. Depken, D. Dulin, The nucleotide addition cycle of the SARS-CoV-2 polymerase. Cell Rep 36, (2021).

47. M. Seifert, P. van Nies, F. S. Papini, J. J. Arnold, M. M. Poranen, C. E. Cameron, M. Depken, D. Dulin, Temperature controlled high-throughput magnetic tweezers show striking difference in activation energies of replicating viral RNA-dependent RNA polymerases. Nucleic acids research 48, 5591–5602 (2020).

48. K. Kwakwa, A. Savell, T. Davies, I. Munro, S. Parrinello, M. A. Purbhoo, C. Dunsby, M. A. Neil, P. M. French, easySTORM: a robust, lower-cost approach to localisation and TIRF microscopy. J Biophotonics 9, 948–957 (2016).

49. D. Axelrod, Total internal reflection fluorescence microscopy in cell biology. Traffic 2, 764–774 (2001).

50. H. Brutzer, F. W. Schwarz, R. Seidel, Scanning evanescent fields using a pointlike light source and a nanomechanical DNA gear. Nano letters 12, 473–478 (2012).

51. C. S. Smith, N. Joseph, B. Rieger, K. A. Lidke, Fast, single-molecule localization that achieves theoretically minimum uncertainty. Nature methods 7, 373–375 (2010).

52. Y. Suzuki, T. Tani, K. Sutoh, S. Kamimura, Imaging of the fluorescence spectrum of a single fluorescent molecule by prism-based spectroscopy. FEBS letters 512, 235–239 (2002).

53. T. Haga, T. Sonehara, T. Fujita, S. Takahashi, Prism-based Spectral Imaging of Four Species of Single-molecule Fluorophores by Using One Excitation Laser. Journal of Fluorescence 23, 591–597 (2013).

54. P. M. Lundquist, C. F. Zhong, P. Zhao, A. B. Tomaney, P. S. Peluso, J. Dixon, B. Bettman, Y. Lacroix, D. P. Kwo, E. McCullough, M. Maxham, K. Hester, P. McNitt, D. M. Grey, C. Henriquez, M. Foquet, S. W. Turner, D. Zaccarin, Parallel confocal detection of single molecules in real time. Optics letters 33, 1026–1028 (2008).

55. J. Eid, A. Fehr, J. Gray, K. Luong, J. Lyle, G. Otto, P. Peluso, D. Rank, P. Baybayan, B. Bettman, A. Bibillo, K. Bjornson, B. Chaudhuri, F. Christians, R. Cicero, S. Clark, R. Dalal, A. deWinter, J. Dixon, M. Foquet, A. Gaertner, P. Hardenbol, C. Heiner, K. Hester, D. Holden, G. Kearns, X. Kong, R. Kuse, Y. Lacroix, S. Lin, P. Lundquist, C. Ma, P. Marks, M. Maxham, D. Murphy, I. Park, T. Pham, M. Phillips, J. Roy, R. Sebra, G. Shen, J. Sorenson, A. Tomaney, K. Travers, M. Trulson, J. Vieceli, J. Wegener, D. Wu, A. Yang, D. Zaccarin, P. Zhao, F. Zhong, J. Korlach, S. Turner, Real-Time DNA Sequencing from Single Polymerase Molecules. *Science (New York*, N.Y 323, 133–138 (2009).

56. J. Jeffet, A. Ionescu, Y. Michaeli, D. Torchinsky, E. Perlson, T. D. Craggs, Y. Ebenstein, Multimodal single-molecule microscopy with continuously controlled spectral resolution. Biophysical Reports 1, (2021).

57. G. Agam, C. Gebhardt, M. Popara, R. Machtel, J. Folz, B. Ambrose, N. Chamachi, S. Y. Chung, T. D. Craggs, M. de Boer, D. Grohmann, T. Ha, A. Hartmann, J. Hendrix, V. Hirschfeld, C. G. Hubner, T. Hugel, D. Kammerer, H. S. Kang, A. N. Kapanidis, G. Krainer, K. Kramm, E. A. Lemke, E. Lerner, E. Margeat, K. Martens, J. Michaelis, J. Mitra, G. G. Moya Munoz, R. B. Quast, N. C. Robb, M. Sattler, M. Schlierf, J. Schneider, T. Schroder, A. Sefer, P. S. Tan, J. Thurn, P. Tinnefeld, J. van Noort, S. Weiss, N. Wendler, N. Zijlstra, A. Barth, C. A. M. Seidel, D. C. Lamb, T. Cordes, Reliability and accuracy of single-molecule FRET studies for characterization of structural dynamics and distances in proteins. Nature methods 20, 523–535 (2023).

58. A. K. Pati, Z. Kilic, M. I. Martin, D. S. Terry, A. Borgia, S. Bar, S. Jockusch, R. Kiselev, R. B. Altman, S. C. Blanchard, Recovering true FRET efficiencies from smFRET investigations requires triplet state mitigation. Nature methods 21, 1222–1230 (2024).

59. F. D. Steffen, R. K. O. Sigel, R. Börner, FRETraj: integrating single-molecule spectroscopy with molecular dynamics. Bioinformatics 37, 3953–3955 (2021).

60. A. N. Kapanidis, N. K. Lee, T. A. Laurence, S. Doose, E. Margeat, S. Weiss, Fluorescence-aided molecule sorting: analysis of structure and interactions by alternating-laser excitation of single molecules. Proceedings of the National Academy of Sciences of the United States of America 101, 8936–8941 (2004).

61. S. Patra, V. Schuabb, I. Kiesel, J.-M. Knop, R. Oliva, R. Winter, Exploring the effects of cosolutes and crowding on the volumetric and kinetic profile of the conformational dynamics of a poly dA loop DNA hairpin: a single-molecule FRET study. Nucleic Acids Research 47, 981–996 (2019).

62. L. E. Baltierra-Jasso, M. J. Morten, L. Laflör, S. D. Quinn, S. W. Magennis, Crowding-induced hybridization of single DNA hairpins. Journal of the American Chemical Society 137, 16020–16023 (2015).

63. D. A. Nicholson, B. Jia, D. J. Nesbitt, Measuring excess heat capacities of deoxyribonucleic acid (DNA) folding at the single-molecule level. The Journal of Physical Chemistry B 125, 9719–9726 (2021).

64. D. A. Nicholson, A. Sengupta, H.-L. Sung, D. J. Nesbitt, Amino acid stabilization of nucleic acid secondary structure: kinetic insights from single-molecule studies. The Journal of Physical Chemistry B 122, 9869–9876 (2018).

65. R. J. Menssen, A. Tokmakoff, Length-dependent melting kinetics of short DNA oligonucleotides using temperature-jump IR spectroscopy. The Journal of Physical Chemistry B 123, 756–767 (2019).

66. A. T. Jonstrup, J. Fredsøe, A. H. Andersen, DNA hairpins as temperature switches, thermometers and ionic detectors. Sensors 13, 5937–5944 (2013).

67. D. Klaue, R. Seidel, Torsional stiffness of single superparamagnetic microspheres in an external magnetic field. Physical review letters 102, 028302 (2009).

68. M. M. van Oene, L. E. Dickinson, F. Pedaci, M. Kober, D. Dulin, J. Lipfert, N. H. Dekker, Biological magnetometry: torque on superparamagnetic beads in magnetic fields. Physical review letters 114, 218301 (2015).

69. I. De Vlaminck, T. Henighan, M. T. van Loenhout, D. R. Burnham, C. Dekker, Magnetic forces and DNA mechanics in multiplexed magnetic tweezers. PloS one 7, e41432 (2012).

70. P. J. Kolbeck, W. Vanderlinden, G. Gemmecker, C. Gebhardt, M. Lehmann, A. Lak, T. Nicolaus, T. Cordes, J. Lipfert, Molecular structure, DNA binding mode, photophysical properties and recommendations for use of SYBR Gold. Nucleic acids research 49, 5143–5158 (2021).

71. F. S. Papini, M. Seifert, D. Dulin, High-yield fabrication of DNA and RNA constructs for single molecule force and torque spectroscopy experiments. Nucleic acids research 47, e144 (2019).

72. P. J. Kolbeck, M. Tišma, B. T. Analikwu, W. Vanderlinden, C. Dekker, J. Lipfert, Supercoiling-dependent DNA binding: quantitative modeling and applications to bulk and single-molecule experiments. Nucleic acids research 52, 59–72 (2024).

73. D. F. Browning, S. J. Busby, Local and global regulation of transcription initiation in bacteria. Nat Rev Microbiol 14, 638–650 (2016).

74. E. F. Ruff, M. T. Record, Jr., I. Artsimovitch, Initial events in bacterial transcription initiation. Biomolecules 5, 1035–1062 (2015).

75. H. Boyaci, J. Chen, R. Jansen, S. A. Darst, E. A. Campbell, Structures of an RNA polymerase promoter melting intermediate elucidate DNA unwinding. Nature 565, 382–385 (2019).

76. R. M. Saecker, A. U. Mueller, B. Malone, J. Chen, W. C. Budell, V. P. Dandey, K. Maruthi, J.H. Mendez, N. Molina, E. T. Eng, Early intermediates in bacterial RNA polymerase promoter melting visualized by time-resolved cryo-electron microscopy. Nature structural & molecular biology 31, 1778–1788 (2024).

77. R. Sreenivasan, I. A. Shkel, M. Chhabra, A. Drennan, S. Heitkamp, H. C. Wang, M. A. Sridevi, D. Plaskon, C. McNerney, K. Callies, C. K. Cimperman, M. T. Record, Jr., Fluorescence-Detected Conformational Changes in Duplex DNA in Open Complex Formation by Escherichia coli RNA Polymerase: Upstream Wrapping and Downstream Bending Precede Clamp Opening and Insertion of the Downstream Duplex. Biochemistry 59, 1565–1581 (2020).

78. A. Revyakin, R. H. Ebright, T. R. Strick, Promoter unwinding and promoter clearance by RNA polymerase: detection by single-molecule DNA nanomanipulation. Proceedings of the National Academy of Sciences of the United States of America 101, 4776–4780 (2004).

79. S. C. Bera, P. P. B. America, S. Maatsola, M. Seifert, E. Ostrofet, J. Cnossen, M. Spermann, F. S. Papini, M. Depken, A. M. Malinen, D. Dulin, Quantitative parameters of bacterial RNA polymerase open-complex formation, stabilization and disruption on a consensus promoter. Nucleic acids research 50, 7511–7528 (2022).

80. L. J. Friedman, J. P. Mumm, J. Gelles, RNA polymerase approaches its promoter without long-range sliding along DNA. Proceedings of the National Academy of Sciences of the United States of America 110, 9740–9745 (2013).

81. F. Wang, S. Redding, I. J. Finkelstein, J. Gorman, D. R. Reichman, E. C. Greene, The promoter-search mechanism of Escherichia coli RNA polymerase is dominated by three-dimensional diffusion. Nature structural & molecular biology 20, 174–181 (2013).

82. A. Mazumder, R. H. Ebright, A. N. Kapanidis, Transcription initiation at a consensus bacterial promoter proceeds via a’bind-unwind-load-and-lock’ mechanism. eLife 10, (2021).

83. D. Dulin, D. L. V. Bauer, A. M. Malinen, J. J. W. Bakermans, M. Kaller, Z. Morichaud, I. Petushkov, M. Depken, K. Brodolin, A. Kulbachinskiy, A. N. Kapanidis, Pausing controls branching between productive and non-productive pathways during initial transcription in bacteria. Nature communications 9, 1478 (2018).

84. L. Yu, J. T. Winkelman, C. Pukhrambam, T. R. Strick, B. E. Nickels, R. H. Ebright, The mechanism of variability in transcription start site selection. eLife 6, (2017).

85. A. M. Malinen, J. Bakermans, E. Aalto-Setälä, M. Blessing, D. L. V. Bauer, O. Parilova, G. A. Belogurov, D. Dulin, A. N. Kapanidis, Real-time single-molecule studies of RNA polymerase– promoter open complex formation reveal substantial heterogeneity along the promoter-opening pathway. Journal of molecular biology, 167383 (2021).

86. J. Chen, C. Chiu, S. Gopalkrishnan, A. Y. Chen, P. D. B. Olinares, R. M. Saecker, J. T. Winkelman, M. F. Maloney, B. T. Chait, W. Ross, R. L. Gourse, E. A. Campbell, S. A. Darst, Stepwise Promoter Melting by Bacterial RNA Polymerase. Mol Cell 78, 275–288 e276 (2020).

87. C. J. Bustamante, Y. R. Chemla, S. Liu, M. D. Wang, Optical tweezers in single-molecule biophysics. Nature Reviews Methods Primers 1, 25 (2021).

88. K. de Zwaan, R. Huo, M. N. F. Hensgens, H. L. Wienecke, M. Tekpınar, H. Geertsema, K. Grußmayer, High-Throughput Single-Molecule Microscopy with Adaptable Spatial Resolution Using Exchangeable Oligonucleotide Labels. ACS nano 19, 13149–13159 (2025).

89. D. Dulin, I. D. Vilfan, B. A. Berghuis, S. Hage, D. H. Bamford, M. M. Poranen, M. Depken, N. H. Dekker, Elongation-Competent Pauses Govern the Fidelity of a Viral RNA-Dependent RNA Polymerase. Cell Rep 10, 983–992 (2015).

90. I. De Vlaminck, T. Henighan, M. T. van Loenhout, I. Pfeiffer, J. Huijts, J. W. Kerssemakers, A. J. Katan, A. van Langen-Suurling, E. van der Drift, C. Wyman, C. Dekker, Highly parallel magnetic tweezers by targeted DNA tethering. Nano letters 11, 5489–5493 (2011).

91. T. Plénat, C. Tardin, P. Rousseau, L. Salomé, High-throughput single-molecule analysis of DNA–protein interactions by tethered particle motion. Nucleic acids research 40, e89–e89 (2012).

92. M. P. DeJong, Y. Bian, J. E. Ortiz-Cárdenas, B. Figueroa, A. Pant, E. Posadas-Barrera, L. Brixi, M. S. Bauer, A. R. Dunn, P. M. Fordyce, Engineering the mechanosensitivity of single DNA molecules via high-throughput microfluidic force spectroscopy. bioRxiv, 2026.2002.2024.707783 (2026).

93. G. Sitters, D. Kamsma, G. Thalhammer, M. Ritsch-Marte, E. J. Peterman, G. J. Wuite, Acoustic force spectroscopy. Nature methods 12, 47–50 (2015).

94. K. C. Neuman, T. Lionnet, J. F. Allemand, Single-molecule micromanipulation techniques. Annual Review of Materials Research 37, 33–67 (2007).

95. F. Stabile, C. Shaheen, S. Leslie, New frontiers in applied biophysics: advancing drug discovery using single-molecule microscopy. Biophysical Reviews 17, 1807–1833 (2025).

96. A. N. Kapanidis, N. K. Lee, T. A. Laurence, S. Doose, E. Margeat, S. Weiss, Fluorescence-aided molecule sorting: analysis of structure and interactions by alternating-laser excitation of single molecules. Proceedings of the National Academy of Sciences 101, 8936–8941 (2004).

97. S. C. Bera, P. P. America, S. Maatsola, M. Seifert, E. Ostrofet, J. Cnossen, M. Spermann, F. S. Papini, M. Depken, A. M. Malinen, Quantitative parameters of bacterial RNA polymerase open-complex formation, stabilization and disruption on a consensus promoter. Nucleic acids research 50, 7511–7528 (2022).

98. V. Svetlov, I. Artsimovitch, “Purification of bacterial RNA polymerase: tools and protocols” in Bacterial transcriptional control: methods and protocols (Springer, 2015), pp. 13–29.

99. D. G. Gibson, L. Young, R.-Y. Chuang, J. C. Venter, C. A. Hutchison, H. O. Smith, Enzymatic assembly of DNA molecules up to several hundred kilobases. Nature methods 6, 343–345 (2009).

100. B. Miroux, J. E. Walker, Over-production of proteins in Escherichia coli: mutant hosts that allow synthesis of some membrane proteins and globular proteins at high levels. Journal of molecular biology 260, 289–298 (1996).

101. S. Quack, D. Dulin, “Surface Functionalization, Nucleic Acid Tether Characterization, and Force Calibration for a Magnetic Tweezers Assay” in Single Molecule Analysis: Methods and Protocols (Springer, 2023), pp. 403–420.

102. V. Nele, M. N. Holme, U. Kauscher, M. R. Thomas, J. J. Doutch, M. M. Stevens, Effect of formulation method, lipid composition, and PEGylation on vesicle lamellarity: a small-angle neutron scattering study. Langmuir 35, 6064–6074 (2019).

103. S. Gao, J. Liang, C. Tan, J. Ma, An oxygen-scavenging system without impact on DNA mechanical properties in single-molecule fluorescence experiments. Nanoscale 17, 3236–3242 (2025).

