## Supplementary Information for "A multimodal, correlative magnetic tweezers–TIRF platform for high-throughput single-molecule interrogations"

M. S. Feiz *et. al.*

This document contains Supplementary Methods, 9 Supplementary Figures, and 1 Supplementary Table.

### S1. Hardware construction and alignment

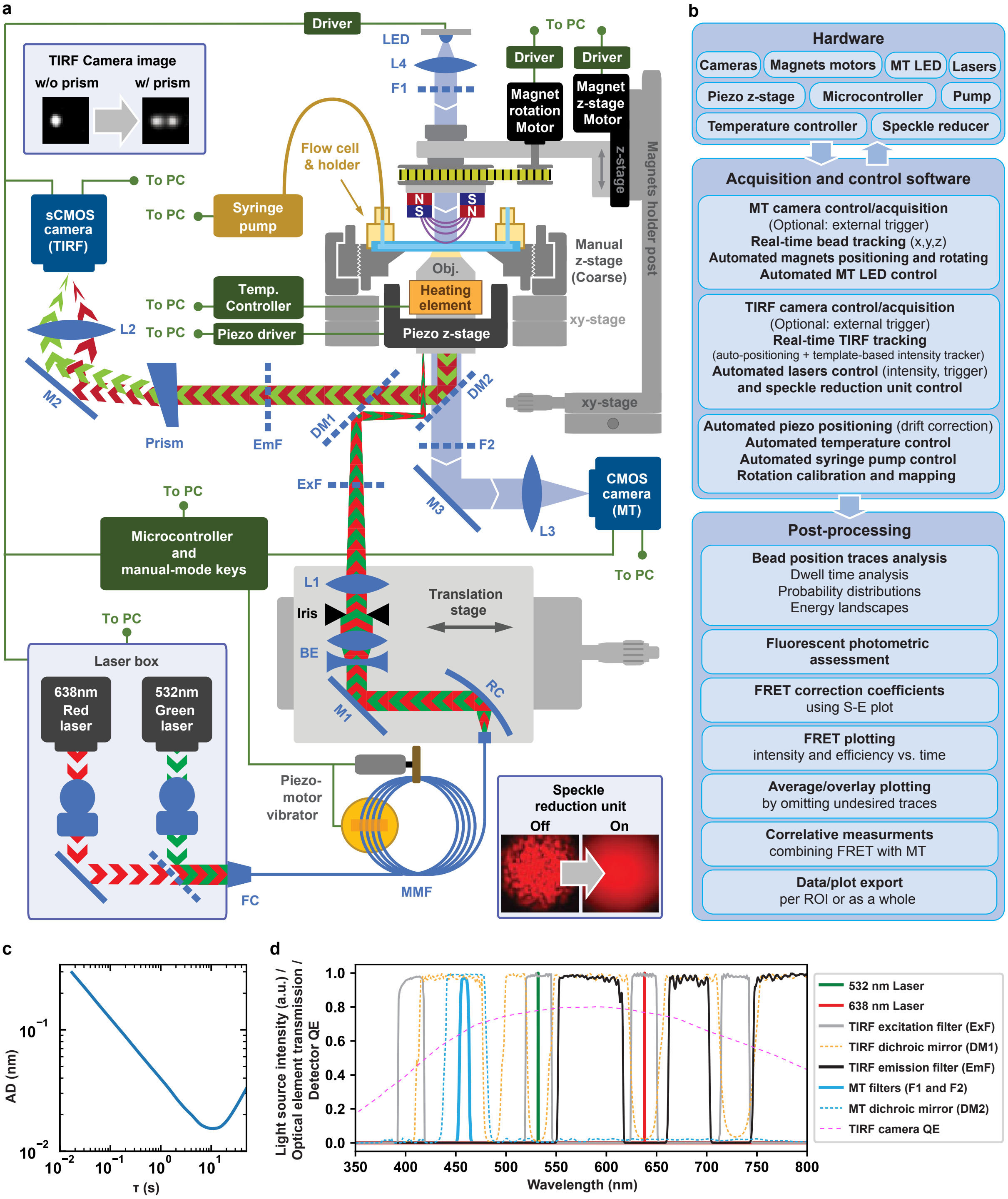


**Fig. S1. High-throughput correlative MT-TIRF assay. (a)** Schematic of the instrument optical paths, hardware and post-processing procedure. The illumination and imaging optical paths are represented for both the MT and TIRF channels. **(b)** Schematic of the sequence of events controlled by the interface. **(c)** Allan deviation (AD) of 3 μm diameter latex reference beads melted on the bottom glass coverslip of the flow chamber after reference bead subtraction. The z-axis position of the reference beads was recorded while using the autofocus as previously described (*1*). **(d)** Transmission spectrum of the optical elements. The quantum efficiency of the TIRF camera is also represented.

**Fig. S1a** shows the schematic for the ray path for the MT-TIRF assay. On the TIRF channel, the laser beam is coupled to the step index multi-mode fiber (MMF) with the help of a laser MMF coupler (FC). The output of the MMF is coupled to the cage system utilizing a reflective collimator (RC), expanded with a beam expander consisting of a lens pair (BE), and cropped by an adjustable iris. Lens L1 focuses the collimated light on the corner of the back focal plane of the Zeiss objective (Obj, 63x, NA = 1.46) to achieve inclined illumination toward the flow chamber, causing total internal reflection (TIR). An excitation filter (ExF) blocks unwanted incoming light. The fluorescence emission from the sample is separated from the excitation source by a dichroic mirror (DM1). The prism decomposes the dyes fluorescence photons to different angles. Finally, the image is formed on the sCMOS camera with a tube lens (L2). On the MT channel, we increased the coherency of the blue LED light with the help of the collimating lens L4 and filter F1. Filter F2 blocks unwanted light from the TIRF channel. The tube lens L3 produces the image on the detector of the CMOS camera. The magnets' rotation and position are controlled by two separate motors (PI Benelux BV). The temperature of the sample is controlled using a heating element (Flexible resistive foil heater, HT10K, Thorlabs) wrapping around the objective. The objective z-position is adjusted using a home-made manual z-stage (~10 µm resolution, ~2 mm range) for coarse adjustments and a piezo z-stage (0.3 nm resolution, 100 µm range, P-726 PIFOC, PI) for fine adjustments.

**Vibrator unit**

To obtain a uniform, top hat-like illumination, we glued a multimode fiber (MMF) bundled in circle to a vibrating piezo element, which scrambles the optical modes of the MMF (*2-4*) (**Fig. S2**). The piezo element (25 mm diameter) was fixed on a 50 mm diameter copper disk and was vibrated at the frequency of ~1.3 kHz provided by an amplified audio signal (Sonitron, PAA-LM4960SQ-02 Amplifier, 24 Vp-p) which was produced by a signal generator (Analog Devices Inc., AD9850) and controlled by a microcontroller unit (Arduino® Due). Optical modes are further scrambled to improve illumination uniformity by inducing higher amplitude vibrations on the MMF using a rotating DC electromotor with a flexible plastic segment hitting the fiber at ~100 Hz (custom-made part). **Fig. S2b** shows how fiber vibration can reduce speckles by mode scrambling in the fiber, including meridional modes, and skew modes.

To measure the speckle contrast, the following equation was used (*5*) :

$$C = \frac{\sigma}{< I >} \quad \text{(S1)},$$

where $\sigma$ and $< I >$ are the standard deviation and the mean intensity of the taken image, respectively, for the entire field of view. Therefore, $C$ reduces with blurring the speckle pattern.

#### S2. Calculations for the TIRF illumination

**Optimal laser beam size**

We used a sCMOS PCO Edge 4.2 LT camera to image the fluorescence photons. The sensor of this camera is a square chip of $2048 \times 2048$ pixels, with 6.5 µm pixel pitch. The total active area is therefore $(13.3 \times 13.3)\, mm^2$, and has a diagonal of 18.8 mm. The field of view (FoV) in the sample space is therefore:

$$\mathrm{D_{FoV}} = \frac{\mathrm{D_{Detector}}}{\text{Objective magnification}} = \frac{18.8}{63} = 0.298\ \mathrm{mm} = 298\ \mathrm{\mu m} \quad \text{(S2)}$$

Where $D_{FoV}$ denotes the diagonal of the sensor image projected on the sample, leading to $(211 \times 211)\, \mu m^2$ FoV, with a pixel size of $103\, nm$ in the sample plane. The excitation beam should cover therefore an area with a diameter of D ~ 300 µm.

The effective focal length of the objective is obtained from:

$$\text{Objective magnification} = \frac{f_{L2}}{f_{Obj}} \quad \text{(S3)},$$

where $f_{Obj}$ and $f_{L2}$ are the objective and tube lens focal lengths respectively. Having $f_{L2} = 165\ mm$ from the provider:

$$f_{Obj} = \frac{f_{L2}}{\text{Objective magnification}} = \frac{165\ \text{mm}}{63} = 2.62\ \text{mm} \quad \text{(S4)}.$$

In **Fig. S1a**, the lens L1 ($f_{L1} = 150\ mm$) and the objective form an infinity imaging lens pair with the magnification of:

$$\text{L1 magnification} = \frac{f_{L1}}{f_{Obj}} = 57.25 \quad \text{(S5)}.$$

Considering the image of the iris is projected on the sample, we can change the sample illumination size by adjusting the iris pupil size. In our case, the diameter of the iris $D_{iris}$ is determined by:

$$D_{iris} = \text{L1 Magnification} \times D_{FoV} = 57.25 \times 298\ \mu\text{m} = 17\ \text{mm} \quad \text{(S6)}.$$

At the MMF output end, we used a reflective collimator (RC, $f_{RC} = 50.8\ \text{mm}$) to collimate the MMF output beam and then a beam expander (BE, composed of an $f = -50\ \text{mm}$ concave lens and an $f = 75\ \text{mm}$ convex lens, resulting in 1.5x magnification). The beam diameter is confined by either the RC acceptance cone angle ($\text{NA} = 0.216,\ \theta_{RC} = 12.47°$) or MMF exit cone angle ($\theta_{MMF} = 12.7°$) and is calculated from the equation below, using $\theta_{RC}$ as the dominant confiner, the beam diameter after reflecting collimator would be:

$$n \sin \theta_{RC} = \frac{D_{Beam}}{2 f_{RC}} \quad \text{(S7)},$$

and therefore:

$$D_{Beam} = 21.94\ \text{mm} \quad \text{(S8)}.$$

After the beam expander (BE), the beam diameter is 1.5x larger, and therefore:

$$D_{Beam\_after\ BE} = 1.5 \times 21.94\ \text{mm} = 32.9\ \text{mm} \quad \text{(S9)}.$$

However, as the beam has Gaussian beam profile, to improve illumination homogeneity (closer to a top-hat profile), the expanded beam must be cropped to 17 mm diameter using the iris.

A translation stage is used to displace the excitation beam parallel to the objective optical axis to adjust its position at the back focal plane of the objective, making it possible to switch between TIRF mode and epi mode. As the incident light at the coverslip-water interface must obey the condition of total internal reflection, the incident ray angle must exceed the critical angle $\theta_c$, i.e. the exciting laser must be focused on the objective back focal plane at a radius larger than $R_c$ from the optical axis. Thus, the excitation beam is confined to an annulus between $R_c$ and the objective back focal plane pupil radius $R_{pupil}$.

The critical angle is calculated from:

$$\sin \theta_c = \frac{n_s}{n_g} \quad \text{(S10)},$$

where $n_s$ is the sample refractive index and $n_g$ is the glass coverslip refractive index (and used for immersion oil refractive index as well). Here, we have:

$$\sin \theta_c = \frac{1.338}{1.515} = 0.883 \quad \text{(S11)}.$$

Thus $R_c$ is:

$$R_c = f_{Obj}\, n_g\, sin\, \theta_c = 2.62\ \text{mm} \times 1.515 \times 0.883 = 3.505\ \text{mm} \quad \text{(S12)}.$$

$R_{pupil}$ is determined from:

$$R_{pupil} = f_{Obj}\, NA = 2.62\ mm \times 1.46 = 3.825\ mm \qquad (S13),$$

where NA is the objective numerical aperture. The annular ring width ($w$) in the objective back focal plane enabling TIR illumination is therefore:

$$w = R_{pupil} - R_c = 320\ \mu m \qquad (S14).$$

In order to stay in a TIR condition, the excitation spot on the back focal plane must be smaller than or equal to $w$. We adjusted the image size of the MMF core at the back focal plane of the objective by tunning RC, BE, and L1 elements. Practically, the MMF exit face was placed at the RC focal point and L1 distance to the objective was adjusted such that in epi-mode, the beam diameter was minimized on a distant screen (in respect to $f_{Obj}$) above the objective. Afterwards, by translating excitation arm (elements on the translation stage in **Fig. S1a**) position parallel to the optical axis to bring the excitation laser beam in TIR, i.e. within the annular region of the back focal plane, a distinct back-reflected spot becomes visible in the illumination path optics.

**W**edge prisms induced spectral dispersion

To decompose the emitted photons from dyes of different wavelengths, a pair of N-BK7 wedge prisms were used. One has a wedge angle of 0° 58' ($\theta_1 = 0.967°$) and the other one has a wedge angle of 7° 41' ($\theta_2 = 7.683°$). We characterized the spatial separation of the photons emitted by Cy3B and ATTO647N, using their emission maxima (570 nm and 665 nm, respectively). As shown in **Fig. S4** the ray trajectory changes twice, at the first prism interface and upon exiting the second prism. Let $\varepsilon_1$ be the incident angle relative to the main optical axis, Snell-Descartes law gives (*6*):

$$\theta_1^i = \theta_1 - \varepsilon_1 \qquad (S15),$$

$$\theta_1^r = sin^{-1}\left(\frac{sin\,\theta_1^i}{n_{prism}}\right) \qquad (S16),$$

$$\theta_2^i = \theta_2 - \theta_1 + \theta_1^r \qquad (S17),$$

$$\theta_2^r = sin^{-1}(n_{prism} sin\,\theta_2^i) \qquad (S18),$$

where the subscripts denote the surface number and the superscripts show incidental ($i$) or refracted ($r$) rays. $\theta$ is measured relative to the surface normal. Referenced to the horizontal axis (main optical axis), this gives:

$$\varepsilon_2 = \theta_2^r - \theta_2 \qquad (S19),$$

$$\varepsilon_2 = sin^{-1}\left(n_{prism} sin\left(\theta_2 - \theta_1 + sin^{-1}\left(\frac{sin(\theta_1 - \varepsilon_1)}{n_{prism}}\right)\right)\right) - \theta_2 \qquad (S20).$$

At the detector plane, under the small-angle approximation, the position of the detected spot relative to the optical axis is:

$$d_{mm} = f_{L2} \times tan\,\varepsilon_2 = 165\ mm \times tan\,\varepsilon_2 \qquad (S21),$$

if the wedge prism is aligned with the sCMOS camera optical axis, the radial distance from the center can be expressed in pixels as:

$$d_{pixel} = d_{mm} \times \frac{\text{detector size}_{pixel}}{\text{detector size}_{mm}} = d_{mm} \times \frac{2048}{13.3} \quad \text{(S22).}$$

**Table S1** shows deviation angles and spot positions for different wavelengths. The wavelengths which are filtered by the emission filter are colored in grey. The calculation was repeated for different incident angles ($\varepsilon_1$) corresponding to different regions in the sample, resulting in changes in spot elongation. However, according to equation S18, the resulting variation is less than a pixel on the detector and hence negligeable, thus we assumed $\varepsilon_1 = 0$.

**Table S1.** Deviation angle and position for different wavelengths after wedge prism

| λ (nm) | Refractive index* | Deviation (Degree) | Spot position deviation (mm) | Spot position deviation (pxl) | Relative spot position (pxl) | Emission peak / Comments |
|---|---|---|---|---|---|---|
| 540 | 1.51893 | 3.5259 | 10.166 | 1566 | -2 | SYBR™ Gold (~537 nm) and orange latex bead (~540 nm) |
| 550 | 1.51844 | 3.5225 | 10.157 | 1564 | 0 | Emission is filtered up to ~550 nm |
| 560 | 1.51797 | 3.5193 | 10.148 | 1563 | 1 | |
| 570 | 1.51753 | 3.5163 | 10.139 | 1561 | 3 | SYTOX™ Orange (~570 nm) and Cy3B (~570 nm) |
| 580 | 1.51711 | 3.5134 | 10.131 | 1560 | 4 | TetraSpeck™(~580 nm, red-orange) |
| 610 | 1.51596 | 3.5056 | 10.108 | 1556 | 8 | |
| 620 | 1.51561 | 3.5032 | 10.101 | 1555 | 9 | Emission is filtered in the range of ~620 nm to ~660 nm |
| 660 | 1.51432 | 3.4943 | 10.075 | 1551 | 13 | |
| 670 | 1.51402 | 3.4923 | 10.070 | 1551 | 13 | ATTO647N (~669 nm) |
| 680 | 1.51373 | 3.4903 | 10.064 | 1550 | 14 | TetraSpeck™ (~680 nm, far-red) |
| 690 | 1.51344 | 3.4884 | 10.058 | 1549 | 15 | |
| 700 | 1.51316 | 3.4865 | 10.053 | 1548 | 16 | Emission is filtered in the range of ~700 nm to ~750 nm |
| 750 | 1.51186 | 3.4776 | 10.027 | 1544 | 20 | |
| 760 | 1.51162 | 3.4759 | 10.022 | 1543 | 21 | |
| 800 | 1.51068 | 3.4695 | 10.004 | 1540 | 24 | |

* For SCHOTT N-BK7 material

To test the imaging system, we first used TetraSpeck™ beads (Invitrogen TetraSpeck Microspheres, 0.1 µm) which have four excitation bands centered at 360, 505, 560, and 660 nm, with corresponding emission maxima at 430, 515, 580, and 680 nm, respectively. When using TetraSpeck™ beads or combination of Cy3B (emission maxima at 571 nm) and ATTO647N (emission maxima at 663 nm) fluorophores, the prism spectrally disperses the emitted light and elongates the spot by ~10 pixels (**Table S1**, column: Relative spot position (pxl), also see **Fig. S4c-e**). A gap is always noticeable in the elongated spot, originating from the multiband dichroic mirror (**Fig. S1d**) reflecting the excitation bandwidth. **Fig. S4c** shows the average intensity profile along the elongated direction, showing the discontinuity in the emission intensity.

The measured separation between the two spots (9.6±0.6 pixels; **Fig. S4cd**) agrees with theoretical predictions (**Table S1**, column: Relative spot position (pxl)), with a dispersion of ~1 pixel over the entire field of view (**Fig. S4de**). Notably, this spectral separation strategy can be readily extended to longer-wavelength emitters, exhibiting an approximately linear relationship between spot separation and emission wavelength over the range of 500–800 nm (**Fig. S4b**).

It is worth noting that the prism displaces the whole image by ~10 mm on the detector relative to the optical axis (**Table S1**, column: Spot position deviation (mm)), which is compensated by a kinematic mirror (M2 in **Fig. S1a**).

**S3. Sample preparation**

Below the structures used in our assays are listed, all the sequences are represented in the 5' to 3' direction, and the colored letters show labeling positions.

*Short duplex DNA assembly for single-molecule FRET experiments*
The oligonucleotides (provided by biomers.net GmbH, Ulm, Germany) consist of 44 bases.

Oligo a: Cy3B labeled oligo, with Cy3B at position 37:
ATTGAGCTGAAAGTGTCGAAGTTGTTTGAGTGTTTGTCTGGATT

Oligo b: The complementary part of the oligo a, 5' biotinylated:
AATCCAGACAAACACTCAAACAACTTCGACACTTTCAGCTCAAT

Oligo c1, c2: ATTO647N labeled oligo, with ATTO647N at position 35 (low-FRET) or 19 (high-FRET), 5' biotinylated:
AATCCAGACAAACACTCAAACAACTTCGACACTTTCAGCTCAAT

Oligo d: The complementary part of the oligo c1 and c2:
ATTGAGCTGAAAGTGTCGAAGTTGTTTGAGTGTTTGTCTGGATT

The combination of oligo a+c1 and oligo a+c2 make the low- and high-FRET populations, respectively. While non-labeled complementary oligos are used to provide A-only and D-only populations.

*DNA FRET hairpin*
The underlined region is the complementary region for self-annealing of the hairpin
Oligo H2: 5' biotinylated, and Cy3B labeled at the 3' end.
TGGCGACGGCAGCGAGGCTTAGCGGCAAAAAAAAAAAAAAAAAAAAAAAAAAAAAAAAGCCGCT

Oligo A2: Atto647N labeled at position 8
GCCTCGCTGCCGTCGCCA

*DNA construct to monitor $RP_O$ dynamics: LacCONS tether sequence*
The construct is amplified via PCR using the pMA-T vector (Thermo Fisher Scientific) according to Stal Papini et al. (*7*) and Bera et al. (*8*). Note that small letters do not align to the template DNA and are used to introduce restriction sites which are underlined.

Digoxigenin handle:
Primer_Dig_fwd:
TCCTCAGCTGAACAAATCTATTATACC
Primer_Dig_rev:
agtgggatccCTGATGCTATAACATGCGCG

Spacer with lacCONS sequence:
Primer_Spacer_fwd:
gcttggatccTTAAGTCGGGACAATAGGGG
Primer_Spacer_rev:
gtacgtgcacAACGCCGAGGATACAC

Biotin handle:
Primer_Bio_fwd:
cacagtgcacTGGCGTACATCTGGATTGAC
Primer_Bio_rev:
GTGATGACCATTTCGGGCG

Complete construct sequence (Dig- (5’) and Bio-(3’) handles are underlined, lacCONS promoter region is bold and italic):

5’-
TCCTCAGCTGAACAAATCTATTATACCAATCGGCTTCAACAATGGTGCTCCACGGCAGGCGCCTGACGAGAGGACCAACACA
CCGAGGAACCTCAGCGCTTCGACATGGCAAAATCCCCCCCTTGCAACTTCTAGAGGAGAAGAGTACTGACTTGAGCGCTCCC
AGCACAACAAGGATAGACTCCTCAGCGTTGCACGTTGGGGATAGCGCTAGCTAACAAAGACGCCTGCTACAACAGGAGTATC
AAACCCGTACAAAGGGAACATCCACACTCCTCAGCTTGGTGAATCGAAGCGCGGCATCAGGATTTCCTTTTGGATACCTGAA
ACAAAGCCCATCGTGGTCCTGTAGACTTGGCACACTCCTCAGCTACACCTGCAGCGCGCGCATGTTATAGCATCAGGCAGTT
TAAGTCGGGACAATAGGGGCCGCAATACACAGTTTACCGCATCCTCAGCCTTGACCTAACTGACAAACTGCCATGGACGACT
AGCCATGCTCTTAGACAGCCCGTCTCATACAGTGATTATGGTCTCGAATTGTCCTCAGCTTG***CAGTGAGCGCAACGCAATAA***
***ATGTGATCTAGATCACATTTTAGGCACCCCAGGCTTGACACTTTATGCTTCGGCTCGTATAATGTGTGGAATTGTGAGAGCG***
***GA***AGGACCTCAGCATTAGAGTTCAATAAGGTCTCCTACCAAGCAACTCAGAGATCTCACAGGCTTAGAAGACCATCAATCTC
CCCTCAGCAGACAGGCCTCCTGTTAAGATGGCAGAGCCCGGTAATCGCACTCAAGACACATAGACTAGTATTCAGGCCTGCT
GGTAATCGCAGGCCTTTTTATTTGGGCGGGCCTCAGCGCATGGCTAACTTGAATTCCTACGTGCGAGGGCAGAAGACTTATC
CGCATTTCGTCTCTTCACCTATCTACTACCCATGCCCCTCAGCCGGAGATTATGTAGGTTGTGAGATGCGGGAGAGGTTCTC
GATCTTCCCGTGGGACGTCAACCTTTCCCTTGATAAAGCATTCCCTCAGCCGCTCGGGTATGGCAGTAAGTACGCCTTCTGA
ATTGTGCTAACCTTCATCCTTATCAAGGCTTGCTGCCAATGATTAGGATTCCTCAGCATTGCCTTGCGACAGACTTCCTACT
CACACTCGCTCACATTGAGCTACTCGATGGGCCATCAGCTTGACCCGCTCTGTAGGGCCTCAGCTCGCGATTACGTGAGTTA
GGGCTCCGGACTGCGCTGTATAGTCGAATCTGATCTCGCCCCAACAACTGCAAACCCCAACTTACCTCAGCTTTAGATAACA
TGATTAGCCGAAGTTGCACGGGGTGCCCACCGTGGACTCCTCCCCGGGTGTCGCTCCTTCATCTGACAATATCCTCAGCGCA
GCCGCTACCACCATCGATTAATACAACGAACGGTGATGTTGTCATAGATTCGGCACATTTCCCTTGTAGGTGTGAAATCACC
CTCAGCTTAGCTTCGCGCCGAAGTCTTATGGCAAAACCGATGGACTATGTTTCGGGTAGCACCAGAAGTCTATAGCACGTGC
ATCCCAACCTCAGCCGTGGCGTGCGTACACCTTAATCACCGCTTCATGCTAAGGTCCTGGCTGCATGCTATGTTGATACGCC
TGCACTGCTCGTAGACCTCAGCAAATATACGAAGCGGGCGGCCTGGCCGGAGCACTACCGCATCGACGCGTATTCGAATACT
GTTAATTGCTCACACATGAGCACCTCAGCAAATAGTAGACCGTCAATTTCAGCCCTCTCTCGTGTATCCTCGGCGTTGTGTG
TCAAATGGCGTACATCTGGATTGACTCTATGACGGCCTCAGCTATTCGCTCCCGCAGCTTGCCAGCACTTTCAGTATCATGG
GGCCCTTGGTTGAATGACTCCTATAACGGACTGCTGATGGGTAGCCTCAGCGGACGACCCAGAATCTATCGGCCGCAGGCAA
GTCCGATTTTTTGCGTTGATTTTTTAATGCAGAATATGCAGCACGAGTTAATGTCCTCAGCTCCGGTATTTGCAAATCGAAT
GGTTGTTGCTCACGAGTCCACCATGCGAGGATATCTTCCTTCTCAAAGTCTGACAGTTCCTCAGCCAGCAAGATATCTGATT
CCAGGCTTTGGCTTTAGCCGCTTCGGTTCATCAGCTCTGATGCCAATCCACGTGGTGTAATTCCTCAGCCCCTCGCCCGAAA
TGGTCATCAC-3’

#### S4. Calibrating the MT spatiotemporal resolution

The system mechanical stability was evaluated using Allan deviation (AD) analysis by tracking surface immobilized 3μm latex beads’ z-position (Latex beads, polystyrene, LB30, Sigma Aldrich) relative to a reference bead position. The Allan deviation of a particle position is defined as (*9*):

$$AD(\tau) = \sqrt{\frac{1}{2}\langle(\bar{z}_{\tau,j+1} - \bar{z}_{\tau,j})^2\rangle} \qquad \text{(S23)}$$

$$\bar{z}_{\tau,j} = \frac{1}{\tau}\int_{\tau(j-0.5)}^{\tau(j+0.5)} z(t)dt \qquad \text{(S24)},$$

where $\tau$ defines both the sampling interval and the averaging time. the AD is one-half the average difference in position between consecutive intervals of duration $\tau$, averaged over all intervals of duration $\tau$.

The corresponding plots for Allan deviation, beads auto-fluorescence and force calibration are shown in **Fig. S1c** (also see (*8*) for more details) and **Fig. S8**.

#### S5. Calibrating TIRF evanescent field excitation depth

We used the MyOne magnetic bead autofluorescence measure the evanescent-field penetration depth. To this end, we calculated the total fluorescence emitted by a spherical bead. We show below that, for beads with radii much larger than the penetration depth, the autofluorescence intensity follows the same exponential decay as observed for a single-point fluorophore.

Assuming that the intensity of a single-point fluorophore located at height $z$ is given by:

$$I(z) = I_0 e^{-z/d} \quad \text{(S25)},$$

where $I_0$ is the intensity at the interface and $d$ is the evanescent-field penetration depth. Consider a bead of radius $R$ whose bottom surface is located at a distance $z_b$ from the interface. The total autofluorescence intensity emitted by the bead can then be estimated as

$$I^b = \int_{z=z_b}^{z=z_b+R} I(z)\, dA \quad \text{(S26)}.$$

where $dA$ represents concentric rings centered around the south pole of the bead, and the integration is performed over the southern hemisphere. Introducing the auxiliary variable $h$, defined as the vertical distance from the bead center, the integral becomes

$$I^b = 2\pi \int_{h=0}^{h=R} I(z_b + R - h)\, hdh = 2\pi I_0 d \left[R - d\left[1 - e^{-R/d}\right]\right] e^{-z_b/d} = I_0^b\, e^{-z_b/d} \quad \text{(S27)},$$

demonstrating an exponential decay with the characteristic length equal to the evanescent field penetration depth, upon varying the bead height $z_b$. Here, $I_0^b$ denotes the autofluorescence intensity when the bead rests on the bottom surface of the flow cell.

We then performed the measurement using a torsionally constrained (coilable) ~3.2 kbp double-stranded RNA (dsRNA) tether (contour length ~ 900 nm) to anchor MyOne magnetic beads in the flow chamber (*10*). Upon laser excitation, the bead exhibited substantial autofluorescence, providing a reporter of the excitation intensity of the evanescent field. At an applied force of 0.3 pN, the introduction of positive supercoils induced plectoneme formation, thereby reducing the end-to-end extension of the tether and progressively moving the magnetic bead closer to the surface (**Fig. S3e**). The resulting change in bead autofluorescence directly reported the local intensity of the evanescent field

**S6. smFRET benchmark analysis**

smFRET benchmark analysis was performed using alternating laser excitation (ALEX) (*11*). We monitored donor and acceptor fluorescence intensities under both donor and acceptor excitation. The field of view contained four populations of fluorescently labeled constructs: donor-only, acceptor-only, low-FRET, and high-FRET species (oligonucleotide sequences provided in the **Supplementary Information**). ALEX enables determination of both the stoichiometry (S) and the FRET efficiency (E) for each construct.

We first estimated the raw FRET efficiency ($\mathrm{E}^{\mathrm{raw}}$) and raw stoichiometry ($S^{\mathrm{raw}}$) (*12-14*):

$$\mathrm{E}^{\mathrm{raw}} = \frac{\mathrm{f_{D|A}}}{\mathrm{f_{D|A}} + \mathrm{f_{D|D}}} \quad \text{(S28)},$$

$$\mathrm{S}^{\mathrm{raw}} = \frac{\mathrm{f_{D|A}} + \mathrm{f_{D|D}}}{\mathrm{f_{D|A}} + \mathrm{f_{D|D}} + \mathrm{f_{A|A}}} \quad \text{(S29)},$$

where D and A denote donor and acceptor fluorophores, respectively, and $\mathrm{f_{x|y}}$ represents the fluorescence intensity detected from fluorophore y upon direct excitation of fluorophore x. **Fig. S5a** shows the S-E distribution calculated using the above equations. However, $\mathrm{f_{D|A}}$ contains contributions beyond FRET-derived photons:

$$\mathrm{f_{D|A}} = \mathrm{I_{leakage}} + \mathrm{I_{dir\ Ex}} + \mathrm{F^{FRET}} \quad \text{(S30)},$$

where $\mathrm{I_{leakage}}$ is the leakage of the donor photons into the acceptor detection area and $\mathrm{I_{dir\ Ex}}$ represents the unwanted direct excitation of the acceptor by the donor laser. The first term is correlated linearly with the donor emission upon donor laser excitation:

$$I_{leakage} = \delta . f_{D|D} \quad \text{(S31)},$$

where δ is coined the leakage constant. As for donor-only population, the two other terms in **(S30)** are zero, we conclude:

$$f_{D|A} = I_{leakage} \quad \text{(for the donor-only population)} \quad \text{(S32).}$$

and therefore:

$$\delta = \frac{E^{raw}_{D-only}}{1 - E^{raw}_{D-only}} \quad \text{(S33)},$$

where $E^{raw}_{D-only}$ represents $E^{raw}$ for the donor-only population. In practice, the δ value is achieved by fitting a 2D-Gaussian on the S-E scatter plot around $E^{raw} = 0$ and $S^{raw} = 1$, and then the fitting mean E value is used for δ estimation.

For the acceptor-only population, there is no donor-related leakage and no FRET as well, hence **(S30)** becomes:

$$f_{D|A} = I_{dir\,Ex} \quad \text{(for the acceptor-only population)} \quad \text{(S34).}$$

The $I_{dir\,Ex}$ is linearly correlated with the acceptor emission upon acceptor laser excitation such as:

$$I_{dir\,Ex} = \alpha . f_{A|A} \quad \text{(S35)},$$

where α is the correlation constant and can be calculated using $S^{raw}$ value for acceptor-only population:

$$\alpha = \frac{S^{raw}_{A-only}}{1 - S^{raw}_{A-only}} \quad \text{(S36)},$$

where $S^{raw}_{A-only}$ represents $S^{raw}$ for acceptor-only population, which -by definition- should have $S^{raw}_{A-only}$ value close to zero. However, in practice S values rarely equal zero in the plotted ranges of the scatter points, instead they are distributed on positive and negative sides of the horizontal line of S = 0. This is because the numerator of S equals the denominator of E, and hence small S values lead to artificially large E values. Therefore, a two-dimensional Gaussian fit to the scatter plot is inappropriate. Instead, the cumulative histogram on the right-hand side of the scatter plot in **Fig. S5a** is used, which includes datapoints outside the plotting window. A Gaussian fit to this distribution, provides $S^{raw}_{A-only}$ for α estimation.

If we apply the abovementioned corrections, we obtain:

$$E^{cc} = \frac{F^{FRET}}{F^{FRET} + f_{D|D}} \quad \text{(S37)}$$

and

$$S^{cc} = \frac{F^{FRET} + f_{D|D}}{F^{FRET} + f_{D|D} + f_{A|A}} \quad \text{(S38)},$$

where

$$F^{FRET} = f_{D|A} - \delta . f_{D|D} - \alpha . f_{A|A} \quad \text{(S39).}$$

Where $E^{cc}$ and $S^{cc}$ are the spectral crosstalk corrected values for FRET efficiency and stoichiometry, respectively, which are represented in **Fig. S5b**.

It is worth noting that in the **Fig. 2b** and **Fig. S5a-b** scatter plots, Gaussian mixture model (GMM) is used for 2D Gaussian fittings. GMM provides a probabilistic framework for describing normally distributed subpopulations within a heterogeneous dataset. Unlike conventional clustering approaches, mixture models do not require prior assignment of data points to specific subpopulations, allowing the model to identify and learn these subpopulations automatically (*15*). In our case, to perform GMM estimation, the number of populations to be fitted is initially determined manually.

Next, we corrected for differences in quantum yield and detection efficiency between the donor and acceptor dyes, which manifest in the S–E plots as low- and high-FRET populations appearing at stoichiometry values different from 0.5. This correction step, which we call gamma correction, is performed by introducing the correction factors γ and β (*16*):

$$\mathrm{E} = \frac{\mathrm{F}^{\mathrm{FRET}}}{\mathrm{F}^{\mathrm{FRET}} + \gamma.\mathrm{f}_{\mathrm{D|D}}} \quad \text{(S40)}$$

And

$$\mathrm{S} = \frac{\mathrm{F}^{\mathrm{FRET}} + \gamma.\mathrm{f}_{\mathrm{D|D}}}{\mathrm{F}^{\mathrm{FRET}} + \gamma.\mathrm{f}_{\mathrm{D|D}} + \mathrm{f}_{\mathrm{A|A}}/\beta} \quad \text{(S41).}$$

γ and β are estimated from a line connecting the two populations in 1/S vs. E dataset and using the following equations:

$$\gamma = \frac{\mathrm{b} - 1}{\mathrm{m} + \mathrm{b} - 1} \quad \text{(S42),}$$

$$\beta = \mathrm{m} + \mathrm{b} - 1 \quad \text{(S43),}$$

where $\mathrm{m}$ and $\mathrm{b}$ are the slope and the intercept of the connecting line, respectively (*13*). **Fig. 2b** shows the corrected ES histogram after gamma correction.

**Data analysis of folding–unfolding dynamic of the short stem hairpin by single-molecule FRET**

Raw traces were averaged over 4 frames. The data was further sorted by Pearson correlation. Only traces that showed an anticorrelated signal for at least 100 frames were taken into account for HMM analysis. The HMM analysis was performed with 2 states and individually fitted using the Python HMMlearn package (version 0.3.3). The last state detected by the HMM was excluded from the analysis as it is unclear whether the dwell time ended because of a change in state or bleaching of one of the two dyes. Dwell time histograms were fitted with a mono-exponential function. To correct for cutoff-induced bias due to photobleaching, we corrected the dwell time distributions according to (*17*):

$$\tau_{off}^{ML} = \bar{t}\left(1 + \frac{T_{cut}N_{cut}}{\bar{t}N_{rec}}\right) \quad \text{(S44),}$$

with $\tau_{off}^{ML}$ as the bias-corrected cutoff, $\bar{t}$ as the mean dwell time, $T_{cut}$ as the experimental cutoff time, $N_{cut}$ as the number of traces that were not bleached by the end of the experiment, and $N_{rec}$ as the total number of recorded traces in the experiment.

The FRET histograms were binned at 0.03 and fitted with a Gaussian distribution. All histograms are normalized to the area of the bars. The apparent activation energies of the folding and unfolding of the DNA hairpin were determined via an Arrhenius fit:

$$k = e^{\frac{-E_a}{RT}} \quad \text{(S45),}$$

with $k$ as the unfolding or folding rate, $E_a$ as the activation energy, $R$ as the gas constant, and $T$ as the temperature.

**S**7. Calibrating the autofluorescence of different magnetic bead brands

We have evaluated magnetic beads form different manufacturers to find beads with the lowest autofluorescence and the best tracking resolution for use in the MT-TIRF assay. Among them, Compel beads (Bangs Lab. Inc., UMC0100, 3.3 µm diameter, streptavidin-coated) and NEB beads (NEB, S1420S, 1 µm diameter, streptavidin-coated) showed the best overall performance, providing sufficient magnetic force, suitable size, and minimal autofluorescence. There are some other beads with low autofluorescence (for example Absolute Mag, 2.57 µm, SMP-UM35, CD-Bioparticles), but not the best options for both red and green illuminations.

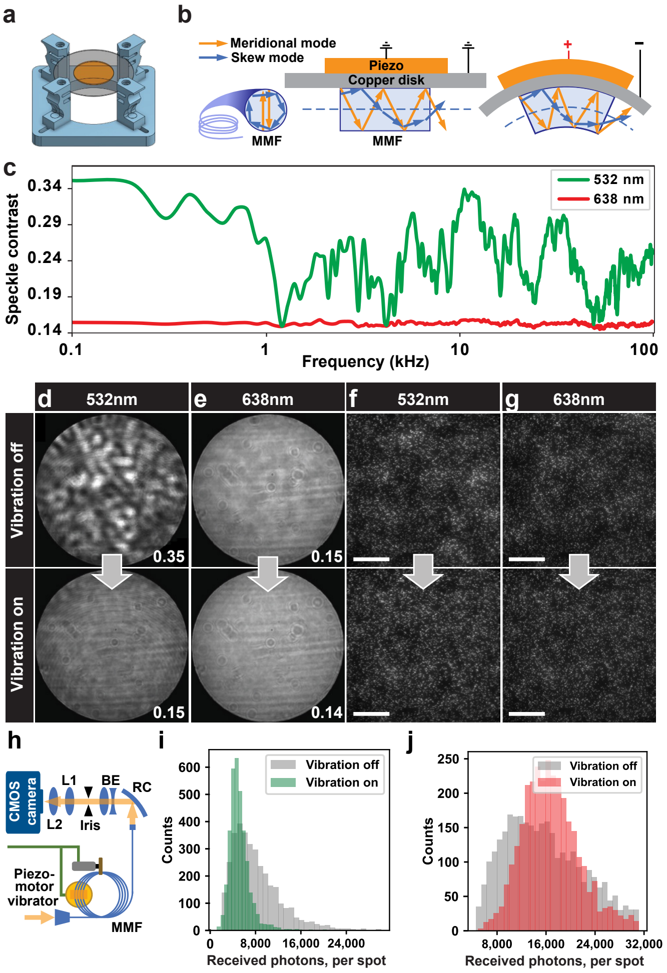


**Fig. S2: Characterization of the speckle reduction unit. (a)** Schematic for the piezo holder. **(b)** Schematic of the effect of MMF bending for different laser mode propagations. **(c)** Speckle contrast as a function of piezo frequencies. **(d, e)** Homogeneity of the laser at the MMF exit face for either 532 nm (d) or 638 nm (e) laser when the piezo vibration is either off or on and set to 1.3 kHz. The speckle contrast value is indicated at the bottom right corner of each image. **(f, g)** Characterization of the laser excitation homogeneity over the field of view of the camera using TetraSpeck microbeads as a fluorescent reporter, excited using either (f) the 532 nm laser (30 W/cm$^2$, 100ms exposure time) or (g) the 638 nm laser (30 W/cm$^2$, 100ms exposure time) with the piezo vibration either off or on. Scale bars show 50 μm. **(h)** Schematic for the speckle imaging arrangement to capture images of (d) and (e). **(i, j)** Histogram of the TetraSpeck bead fluorescence intensity when the piezo vibrator is either off (grey) or on (green and red) when using the 532 nm and 638 nm lasers, respectively.

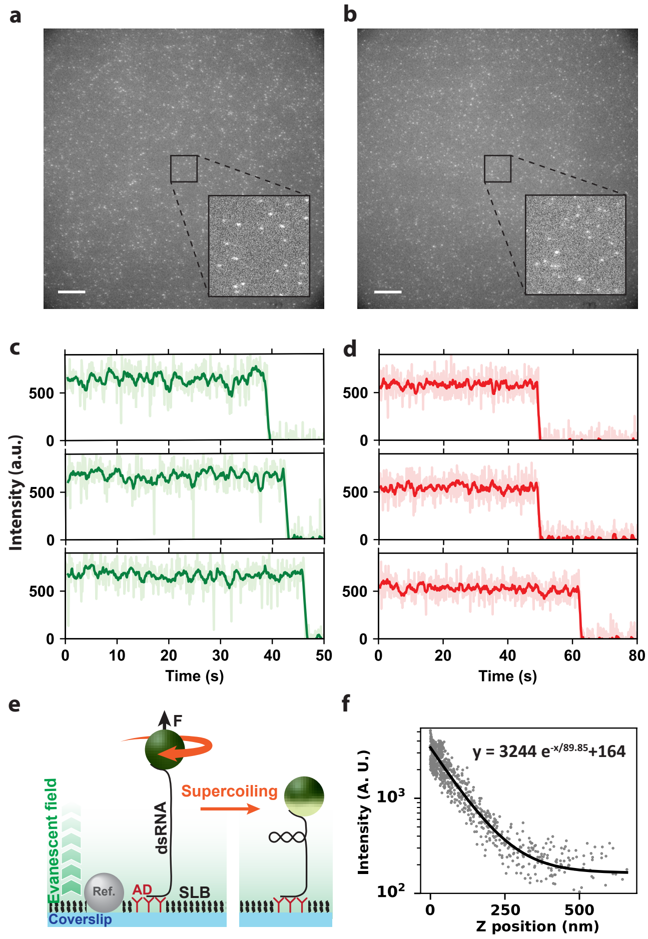


**Fig. S3: TIRF microscope characterization. (a, b)** Field of view with (a) Cy3B labeled oligos and (b) ATTO647N labeled oligos attached to the coverslip surface, respectively excited by 532 nm and 638 nm lasers with 100 W/cm$^2$ intensity and 100 ms exposure time. Insets: zoom-in of the field of view. The scale bars represent 20 µm. **(c, d)** Examples of Cy3B (c) and ATTO647N (d) single dye molecule intensity trajectory versus time, respectively, acquired as described in (a, b). The dark colored traces show the 10-frame boxcar average. **(e)** Schematic of the evanescent field penetration depth. Positively rotating the magnetic bead tethered by a coilable dsRNA construct results in plectonemes formation, driving the magnetic bead downward into the evanescent field, and increasing the magnetic bead autofluorescence intensity. **(f)** Autofluorescence intensity of MyOne beads as a function of the magnetic bead vertical position for the experiment performed as described in (e). The inserted formula is an exponential fit, with a ~90 nm penetration depth.

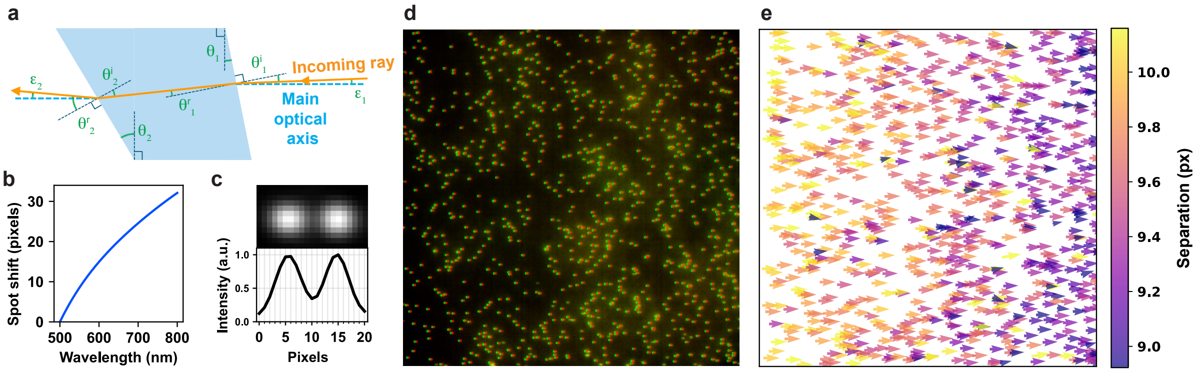


**Fig. S4. Optical elements characteristics. (a)** Beam decomposition through a wedge prism. Only one ray is represented for simplicity. **(b)** Spot shift by the wedge prism as a function of the wavelength, relative to 500 nm wavelength. **(c)** Image and profile of the average from all TetraSpeck fluorescent beads in the field of view obtained when illuminating the sample with the 532 nm and 638 nm lasers. **(d)** Field of view of fluorescent beads illuminated by both green and red lasers, overlapped. **(e)** Displacement vector field showing the detected green-red spots separation over the entire field of view represented in (d).

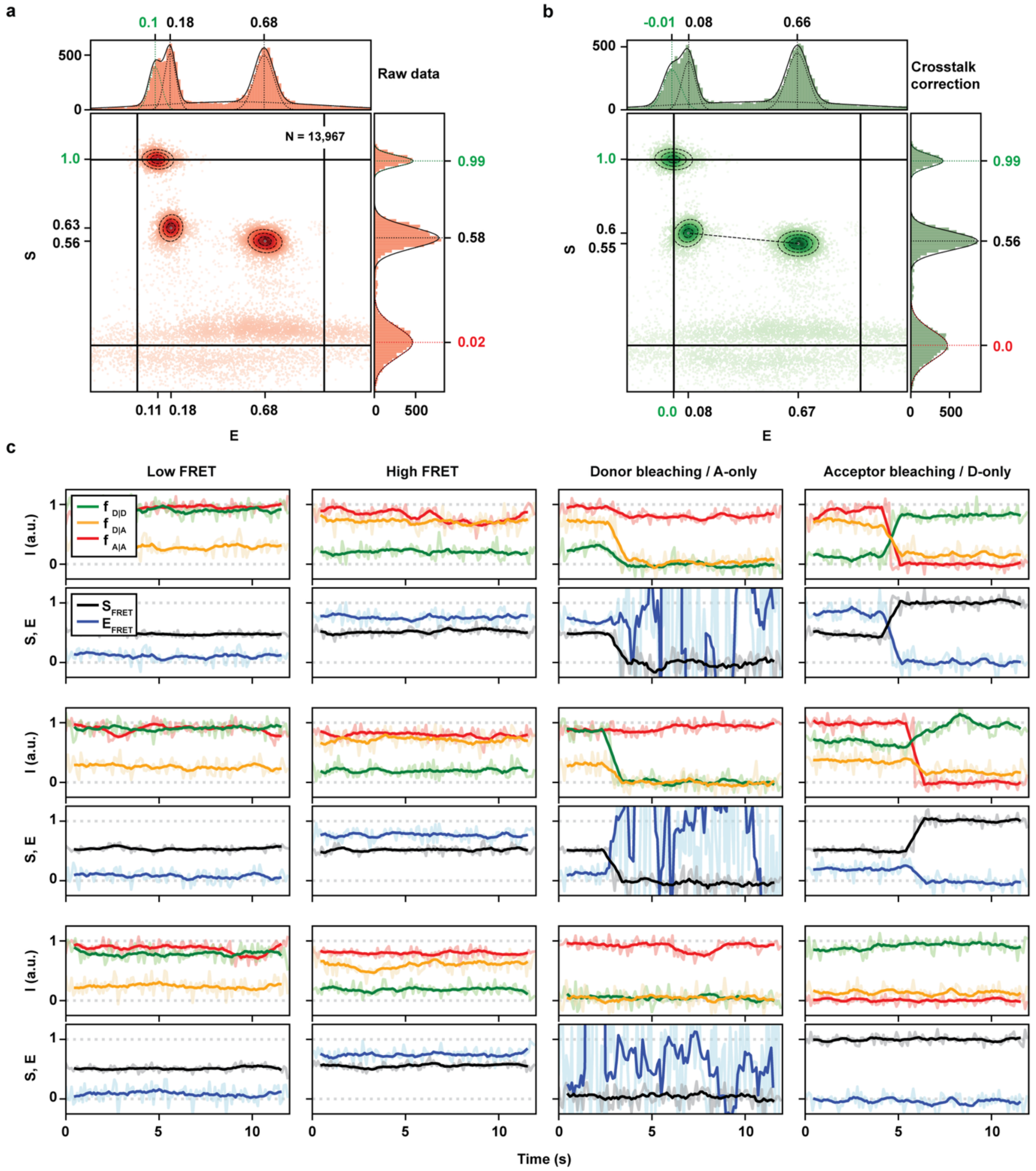


**Fig. S5. TIRF calibration and assessment using high- and low-FRET duplex oligo samples. (a)** Background corrected stoichiometry-efficiency (S-E) plot for the raw data acquired as in Fig. 2a. **(b)** Spectral crosstalk corrected S-E plot. In the panels (a) and (b), the dashed oval lines represent 2D Gaussian fits. E and S histogram are represented on the top and right axis respectively, with 1D-Gaussian fits (solid lines). The values indicated in green and red correspond to the D-only and A-only populations, respectively. **(c)** Fluorescent intensity, FRET efficiency and stoichiometry of the donor and acceptor dyes. From left to right: the first column shows traces for low-FRET pairs. The second column shows high-FRET pairs. The third and last columns show high- and low-FRET traces followed by bleaching of either the donor or the acceptor. The last row of these two columns shows acceptor-only (S~0) and donor-only (S~1) traces. All the traces were acquired at 1 Hz in ALEX mode (light color) and 5-frame boxcar averaged (dark color).

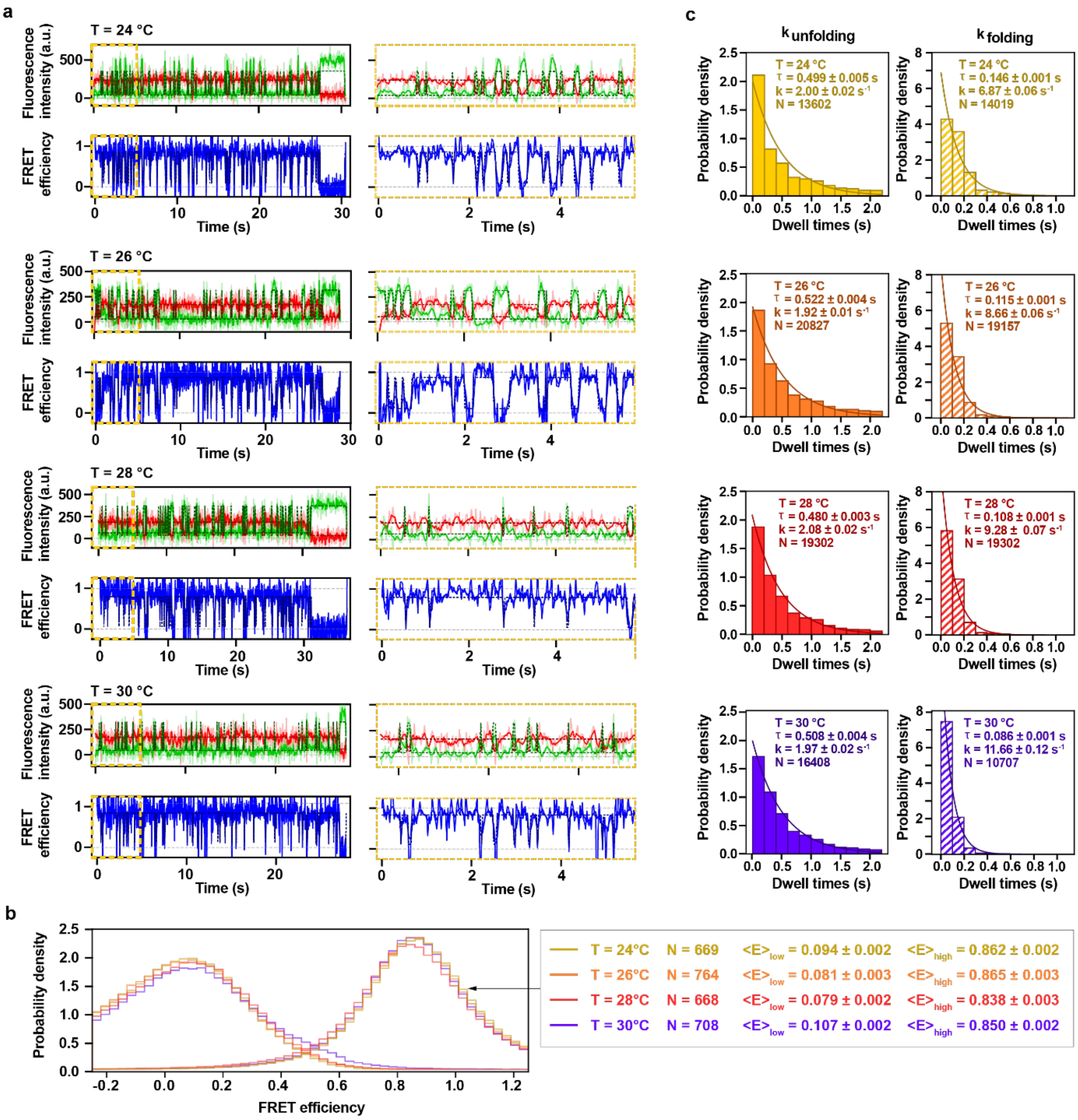


**Fig. S6. DNA hairpin folding and unfolding dynamics as a function of temperature. (a)** smFRET traces of Cy3B and Atto647N labeled DNA hairpin at 24°C, 26°C, 28°C, and 30°C. Anticorrelated fluorescence intensity dynamics of the donor (green) and acceptor (red) dyes, and the resulting FRET efficiency (blue). Light color and dark color represent raw and 4-frame averaged data, respectively. The dashed line is a HMM fit to the raw data. A zoom-in of the first 5 s of each trace is displayed to the right of the full trace (yellow dashed rectangle). **(b)** FRET efficiency distributions. Median FRET efficiencies and error estimation from Gaussian fits, as well as the number of traces n are indicated for the high and low FRET states at 24°C (yellow), 26°C (orange), 28°C (red), and 30°C (purple). **(c)** Dwell time histograms show the unfolding (solid bars) and folding (hatched bars) rates for the same conditions as in (a) and (b). A mono-exponential fit is sketched as a solid line and the mean dwell time ($\tau$) ± standard error of the mean, the reaction rate ($k_{unfolding}$ and $k_{folding}$, respectively) and the number of state detection N are given in the plot.

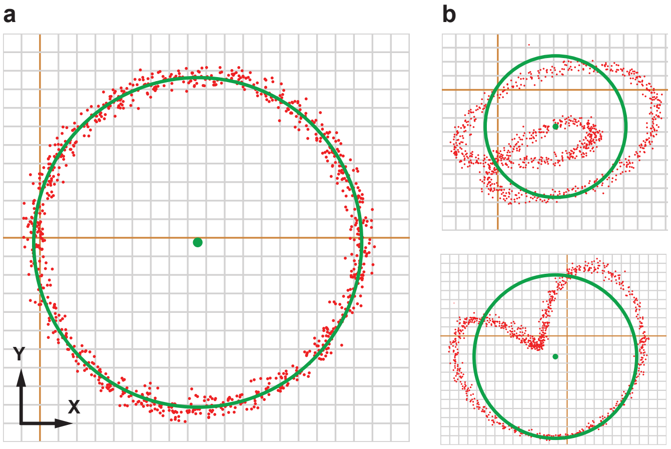


**Fig. S7: Examples of magnetic bead rotation. (a)** When the magnetic field effect is axially symmetric over magnets rotation. **(b)** When the magnetic field configuration results in a non-strictly vertical tether.

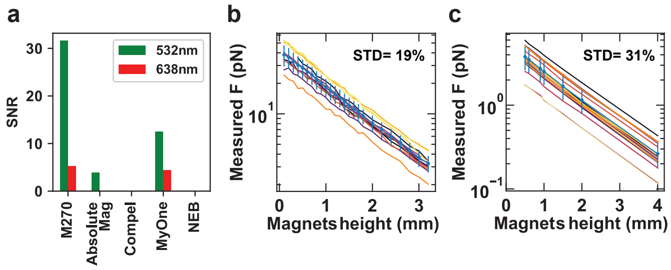


**Fig. S8. Fluorescent background and force calibration of commercial magnetic beads. (a)** Signal-to-noise ratio (SNR) for the auto-fluorescence emission of different types of magnetic beads when illuminated with ~100 W/cm$^2$ of either 532 nm (green) or 638 nm (red) laser. **(b, c)** Magnetic tweezers force-calibration performed with either (b) Compel (N=12) or (c) NEB (N=17) magnetic beads. The blue line is the average force estimation, and the error bars represent one standard deviation.

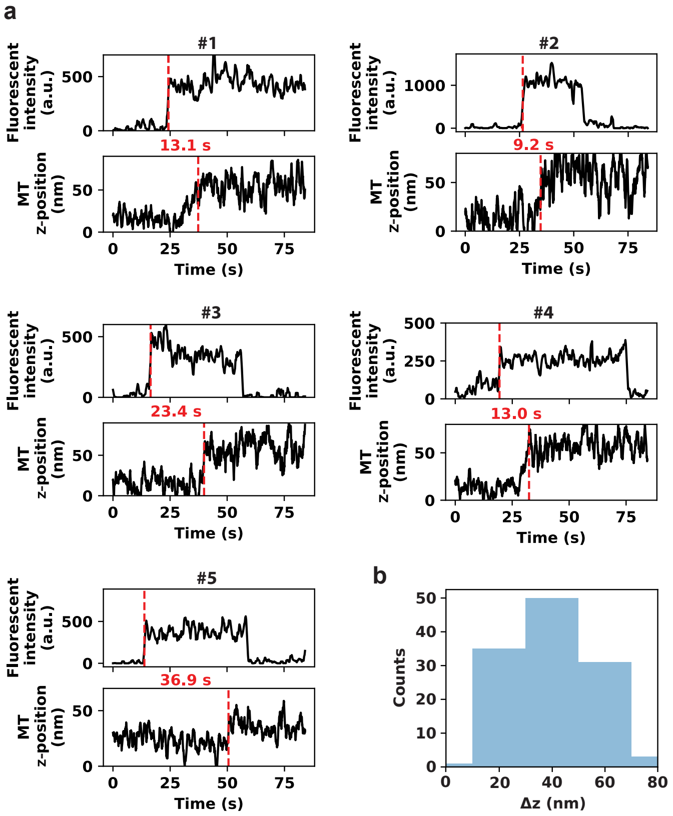


**Fig. S9. Examples of correlative MT-TIRF experimental traces of the open complex formation. (a)** The traces are boxcar averaged with a smoothing window of 1 sec for both channels. The red dashed lines indicate the time at which either the labeled protein binds to the promoter and appears in the TIRF channel, or the transcription bubble opens, changing the tether extension and the magnetic bead position in the MT channel. The red numbers indicate the time delay between the red dashed lines. **(b)** Histogram of the tether extension change from transcription bubble opening during $RP_O$ formation.

#### References


1. S. C. Bera, M. Seifert, R. N. Kirchdoerfer, P. van Nies, Y. Wubulikasimu, S. Quack, F. S. Papini, J. J. Arnold, B. Canard, C. E. Cameron, The nucleotide addition cycle of the SARS-CoV-2 polymerase. *Cell Reports* **36**, (2021).
2. W.-S. Ha, S.-J. Lee, K.-H. Oh, Y.-M. Jung, J.-K. Kim, Speckle reduction in near-field image of multimode fiber with a piezoelectric transducer. *Journal of the Optical Society of Korea* **12**, 126-130 (2008).
3. W. Ha, S. Lee, Y. Jung, J. K. Kim, K. Oh, Acousto-optic control of speckle contrast in multimode fibers with a cylindrical piezoelectric transducer oscillating in the radial direction. *Optics express* **17**, 17536-17546 (2009).
4. D. S. Mehta, D. N. Naik, R. K. Singh, M. Takeda, Laser speckle reduction by multimode optical fiber bundle with combined temporal, spatial, and angular diversity. *Applied optics* **51**, 1894-1904 (2012).
5. J. W. Goodman, *Speckle phenomena in optics: theory and applications* (Roberts and Company Publishers, 2007).
6. E. Hecht, *Optics* (Pearson Education India, 2012).
7. F. S. Papini, M. Seifert, D. Dulin, High-yield fabrication of DNA and RNA constructs for single molecule force and torque spectroscopy experiments. *Nucleic acids research* **47**, e144-e144 (2019).
8. S. C. Bera, P. P. America, S. Maatsola, M. Seifert, E. Ostrofet, J. Cnossen, M. Spermann, F. S. Papini, M. Depken, A. M. Malinen, Quantitative parameters of bacterial RNA polymerase open-complex formation, stabilization and disruption on a consensus promoter. *Nucleic acids research* **50**, 7511-7528 (2022).
9. D. Dulin, "An introduction to magnetic tweezers" in *Single Molecule Analysis: Methods and Protocols* (Springer, 2023), pp. 375-401.
10. F. S. Papini, M. Seifert, D. Dulin, High-yield fabrication of DNA and RNA constructs for single molecule force and torque spectroscopy experiments. *Nucleic acids research* **47**, e144 (2019).
11. A. N. Kapanidis, N. K. Lee, T. A. Laurence, S. Doose, E. Margeat, S. Weiss, Fluorescence-aided molecule sorting: analysis of structure and interactions by alternating-laser excitation of single molecules. *Proceedings of the National Academy of Sciences* **101**, 8936-8941 (2004).
12. J. Hohlbein, T. D. Craggs, T. Cordes, Alternating-laser excitation: single-molecule FRET and beyond. *Chem Soc Rev* **43**, 1156-1171 (2014).
13. N. K. Lee, A. N. Kapanidis, Y. Wang, X. Michalet, J. Mukhopadhyay, R. H. Ebright, S. Weiss, Accurate FRET measurements within single diffusing biomolecules using alternating-laser excitation. *Biophysical journal* **88**, 2939-2953 (2005).
14. E. Lerner, T. Cordes, A. Ingargiola, Y. Alhadid, S. Chung, X. Michalet, S. Weiss, Toward dynamic structural biology: Two decades of single-molecule Forster resonance energy transfer. *Science (New York, N.Y* **359**, (2018).
15. C. M. Bishop, N. M. Nasrabadi, *Pattern recognition and machine learning* (Springer, 2006), vol. 4.
16. N. K. Lee, A. N. Kapanidis, Y. Wang, X. Michalet, J. Mukhopadhyay, R. H. Ebright, S. Weiss, Accurate FRET measurements within single diffusing biomolecules using alternating-laser excitation. *Biophysical journal* **88**, 2939-2953 (2005).
17. B. Eslami-Mosallam, I. Katechis, M. Depken, "Fitting in the age of single-molecule experiments: a guide to maximum-likelihood estimation and its advantages" in *Biophysics of RNA-Protein Interactions: A Mechanistic View* (Springer, 2019), pp. 85-105.
